# Probing the biogenesis pathway and dynamics of thylakoid membranes

**DOI:** 10.1101/2021.05.05.442711

**Authors:** Tuomas Huokko, Tao Ni, Gregory F. Dykes, Deborah M. Simpson, Philip Brownridge, Fabian D. Conradi, Robert J. Beynon, Peter J. Nixon, Conrad W. Mullineaux, Peijun Zhang, Lu-Ning Liu

## Abstract

How thylakoid membranes are generated to form the metabolically active membrane network and how thylakoid membranes orchestrate the insertion and localization of protein complexes for efficient electron flux remain elusive. Here, we develop a method to modulate thylakoid biogenesis in the rod-shaped cyanobacterium *Synechococcus elongatus* PCC 7942 by modulating light intensity during cell growth, and probe the spatial-temporal stepwise biogenesis process of thylakoid membranes in cells. Our results reveal that the plasma membrane and regularly arranged concentric thylakoid layers have no physical connections. The newly synthesized thylakoid membrane fragments emerge between the plasma membrane and pre-existing thylakoids. Photosystem I monomers appear in the thylakoid membranes earlier than other mature photosystem assemblies, followed by generation of Photosystem I trimers and Photosystem II complexes. Redistribution of photosynthetic complexes during thylakoid biogenesis ensures establishment of the spatial organization of the functional thylakoid network. This study provides insights into the dynamic biogenesis process and maturation of the functional photosynthetic machinery.

## Introduction

Oxygenic photosynthesis produces energy and oxygen for life on Earth and is arguably the most important biological process. In phototrophs, such as cyanobacteria, algae, and plants, photosynthetic light reactions are performed by a set of light-harvesting complexes, reaction center protein complexes, as well as other electron transport complexes and cofactors, which are accommodated in specialized intracellular membranes, termed thylakoid membranes^1^. Unlike the thylakoids in plant chloroplasts, cyanobacterial thylakoid membranes do not form tightly appressed grana stacks; they sit between the plasma membrane and the central cytoplasm, leading to intricate cellular compartmentalization and providing the site for both photosynthetic and respiratory electron transport chains^2,3^. The photosynthetic electron transport chain comprises four membrane-bound protein complexes: Photosystem I (PSI), Photosystem II (PSII), Cytochrome (Cyt) *b*_6_*f*, and ATP synthase (ATPase), along with the electron carriers plastoquinone and plastocyanin. The respiratory electron transport chain involves type-I NAD(P)H dehydrogenase-like complex (NDH-1), succinate dehydrogenase, and terminal oxidases^4–7^. The interplay of these electron transport complexes and molecules ensures the physiological direction of electron flows and the required stoichiometry of ATP/NADPH synthesis in the cell^8^. An outstanding fundamental question is how the thylakoid membranes and photosynthetic machinery are constructed and maintain functionality. Despite considerable efforts in determining the atomic structures and spectroscopic properties of individual thylakoid protein complexes, we are only beginning to understand the assembly and membrane locations of photosynthetic complexes, as well as the spatial organization of thylakoids in cyanobacteria. For example, early genetic and biochemical analysis have indicated that the multi-subunit PSII and PSI complexes undergo sophisticated stepwise assembly processes that are finely regulated in time and space^9,10^. Recent atomic force microscopy (AFM) studies have revealed the native organization of different photosynthetic complexes in thylakoids of the cyanobacterium *Synechococcus elongatus* PCC 7942 (*Synechococcus*)^11,12^. The photosynthetic complexes are densely packed in the thylakoid lipid layer, which facilitates the association of functionally coordinated complexes and electron flow. Likewise, the photosynthetic assemblies exhibit high heterogeneity in structure and membrane arrangement, highlighting their structural and assembly variations *in situ*^11^.

Little is known about where the biogenesis of thylakoid membranes originates in cyanobacterial cells and whether there are physical connections or fusions between the intracytoplasmic thylakoid membranes and the plasma membrane. Electron microscopy (EM) on the cyanobacterium *Synechocystis* sp. PCC 6803 (*Synechocystis*) have shown that thylakoid membranes converge at some sites close to the plasma membrane^13,14^. Recently, cryo-electron tomography (cryo-ET) of *Synechocystis* cells did not identify any direct fusion of the plasma membrane and thylakoid membranes^15^; instead, cells possess so-called “thylapse” contacts where thylakoid convergence membranes form tight connections with the plasma membrane^15^. At the convergence regions, distinct layers of thylakoids interconnect with each other and form a contiguous membrane network, which potentially facilitates diffusion of constituents between adjacent thylakoid lumens and electron fluxes throughout the thylakoid network^16,17^. While these studies provided static pictures about the assembly of photosynthetic complexes and the organization of thylakoid membranes, a systematic characterization of the molecular processes concerning the origin and development of thylakoid membranes remains to be established for a complete understanding of the pathways and dynamics of thylakoid biogenesis.

Cyanobacterial thylakoid membranes are truly plastic and adapt rapidly to changes in the growth conditions. Previous studies have revealed that the thylakoid content could be reduced by dark acclimation or nitrogen starvation and could be recovered by exposing cells to light or nitrate^18–20^. The physiological modulations of thylakoid formation, protein composition and organization provide the foundation for exploring in detail the biogenesis process and functions of photosynthetic apparatus in thylakoid membranes. Here, we developed a method to modulate *in vivo* thylakoid biogenesis of *Synechococcus*, by regulating light intensity during cell growth. Unlike the irregular *Synechocystis* thylakoid organization, the thylakoid membranes in the rod-shaped *Synechococcus* cells form regular concentric cylinders aligned along the plasma membrane, suitable for quantitative analysis^8,12^. Using thin-section EM, cryo-ET of cell lamella produced by cryo-focused ion beam (cryo-FIB), live-cell confocal microscopy, proteomics, and biochemical approaches, we probed the initial sites of thylakoid biogenesis and the stepwise membrane integration, assembly, and dynamic localization of photosynthetic complexes during thylakoid biogenesis. The results enabled us to delineate the biogenesis pathway of cyanobacterial thylakoid membranes, which may inform strategies for engineering functional photosynthetic systems to underpin bioenergy production.

## Results

### Manipulation of thylakoid membrane biogenesis by varying light intensity

To control the thylakoid membrane biogenesis, *Synechococcus* cells were first subjected to high light (HL, 300 µmol photons m^−2^ s^−1^) for 14 days to greatly reduce the content of intracellular thylakoids compared with the cells under growth light (GL, 40 µmol photons m^−2^ s^−1^), as indicated by the reduction of pigment content per cell (Supplementary Fig. 1). Thin-section electron microscopy (EM) revealed that unlike the GL-grown cells that possess generally 3-4 layers of mostly intact, concentric thylakoid membranes (Fig. 1a, 1b, Supplementary Fig. 2), the number of thylakoid layers in the HL-adapted cells was greatly reduced. There is typically one continuous thylakoid layer that was accompanied occasionally by a thylakoid membrane fragment close to the plasma membrane (Fig. 1a). The distance between the inner and peripheral thylakoid layers (109.2 ± 1.6 nm, *n* = 50) was ~39% larger than the space between thylakoid layers in the GL-grown cells (78.4 ± 3.4 nm, *n* = 50) (Fig. 1c). The HL-adapted cells were then transferred to low light (LL, 20 µmol photons m^−2^ s^−1^) to promote the development of thylakoid membranes. The changes in pigment content and thylakoid membrane ultrastructures were assayed at distinct time points during LL adaptation (Fig. 1, Supplementary Figs. 1 and 2). In the cells after one-day LL adaptation (LL day 1), one additional thylakoid layer appeared on the same longitudinal side of the cell where the single membrane fragment was observed, giving rise to an asymmetrical distribution of thylakoid membranes (Fig. 1a, 1b). The distance between thylakoid layers (110.6 ± 5.5 nm, *n* = 50) remained relatively consistent with that of HL-adapted cells (Fig. 1c). In LL day 2, the existing thylakoid fragments expanded, generating fascicles of lamellar sheets or spiral-like thylakoid membranes lining the plasma membrane in the cell, reducing the asymmetric distribution of thylakoids (Fig. 1a, 1b). After LL day 2, the content and organization of thylakoid membranes become closer to those of the GL-grown cells; on LL day 5 the cells exhibited a GL-like phenotype (Fig. 1, Supplementary Fig. 1). Thus, by regulating light intensity and duration, we could specifically control thylakoid expansion to characterize the membrane construction process.

**Figure 1.**
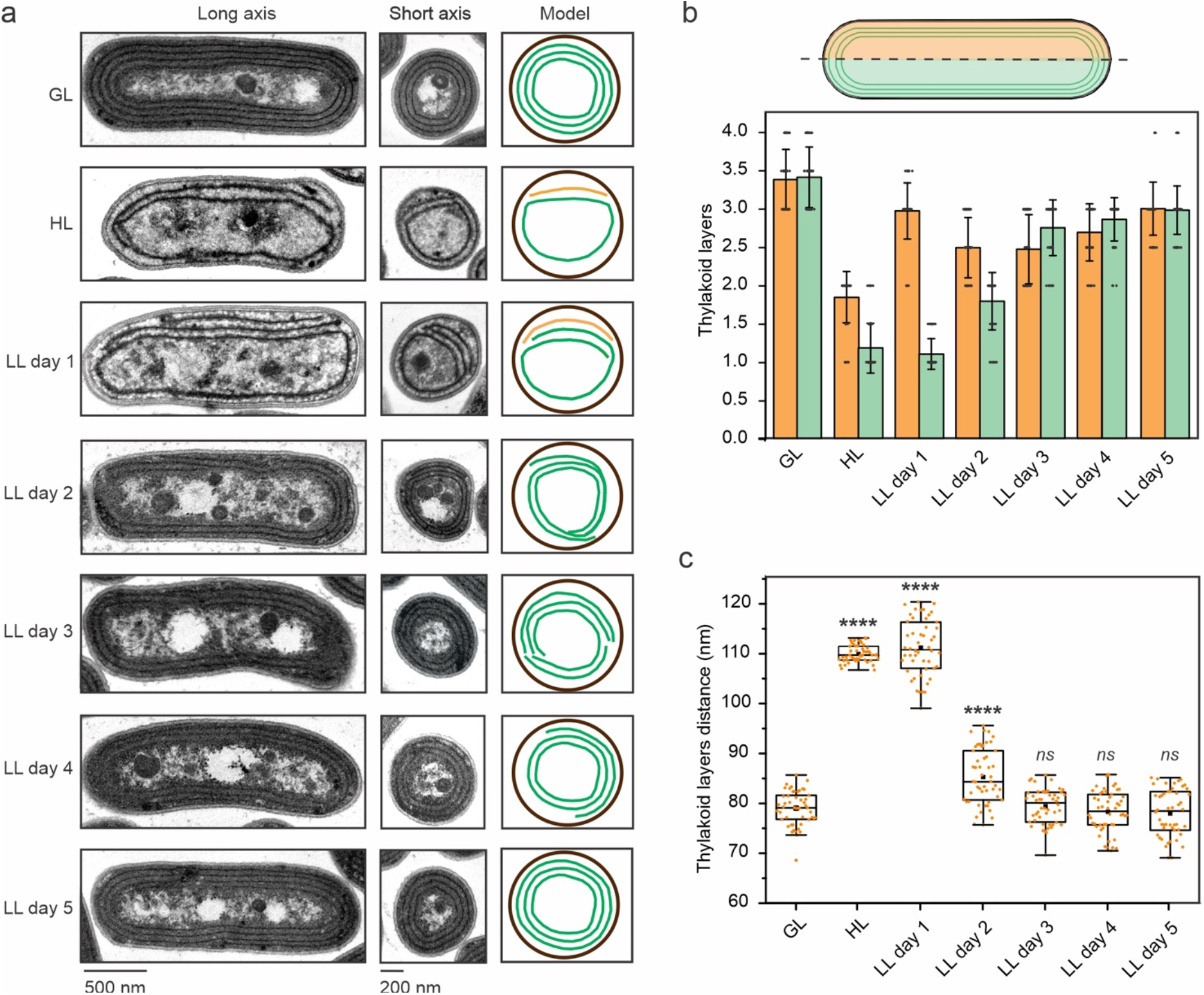
The ultrastructures of thylakoid membranes during its light-regulated biogenesis in *Synechococcus*. Cells were grown under growth light (GL), high light (HL), and HL-grown cells were transferred to low light (LL) conditions for 5 days. **a**, Representative transmission electron microscope (TEM) images of cells grown under GL, HL, and HL-grown cells transferred to LL for 5 days. In the thylakoid elongation models, existing membrane parts are in green and new ones in orange during the initial steps of the biogenesis process. Representative TEM images were derived from at least three biologically independent preparations with similar results. **b**, Number of thylakoid membrane layers on both longitudinal sides of cells. Values are means ± SD; *n* = 50 cells in total from 3 biologically independent experiments **c**, Distance between thylakoid membrane layers. *n* = 50 cells from 3 biologically independent experiments for each condition. Asterisks indicate the statistically significant differences compared to GL cells. Box plots display the median (line), the average (filled square), the interquartile range (box), and the whiskers (extending 1.5 times the interquartile range). For HL *p* = 2.92 ⨯ 10^−77^, for LL day 1 *p* = 5.89 ⨯ 10^−57^ and for LL day 2 *p* = 9.42 ⨯ 10^−10^. *ns*, not significant. Statistical analysis was performed using two-sided two-sample t-Test.

### *In situ* organization of thylakoid membranes in *Synechococcus*

To dissect in-depth the thylakoid membrane organization at the early stage of biogenesis, we applied *in situ* cryo-ET to image the *Synechococcus* cells grown under HL and LL day 1. The HL-induced decrease in the number of thylakoid layers was confirmed by cryo-ET (Fig. 2a, 2b). No apparent connection or extended continuity between the plasma and thylakoid membranes was detected in the *Synechococcus* cells (Fig. 2, Supplementary Fig. 3, 4, Supplementary Movie 1, 2), in agreement with earlier studies from other cyanobacterial species^14,15,21^. Alternatively, the tips of thylakoid extension appeared in close vicinity of the plasma membrane (Fig. 2e, 2f, 2i, 2j, Supplementary Movie 3), resembling the “thylapse” domains as reported in *Synechocystis*^15^. Likewise, small membrane segments were visualized near the thylakoid membrane “breakages” close to the plasma membrane (Supplementary Fig. 5, Supplementary Movies 4, 5). These segments were connected with the thylakoid layer and approached the plasma membrane without direct fusion, probably representing the convergence zones of thylakoid membranes. Similar thylakoid domains have also been observed in *Synechocystis*^15^.

**Figure 2.**
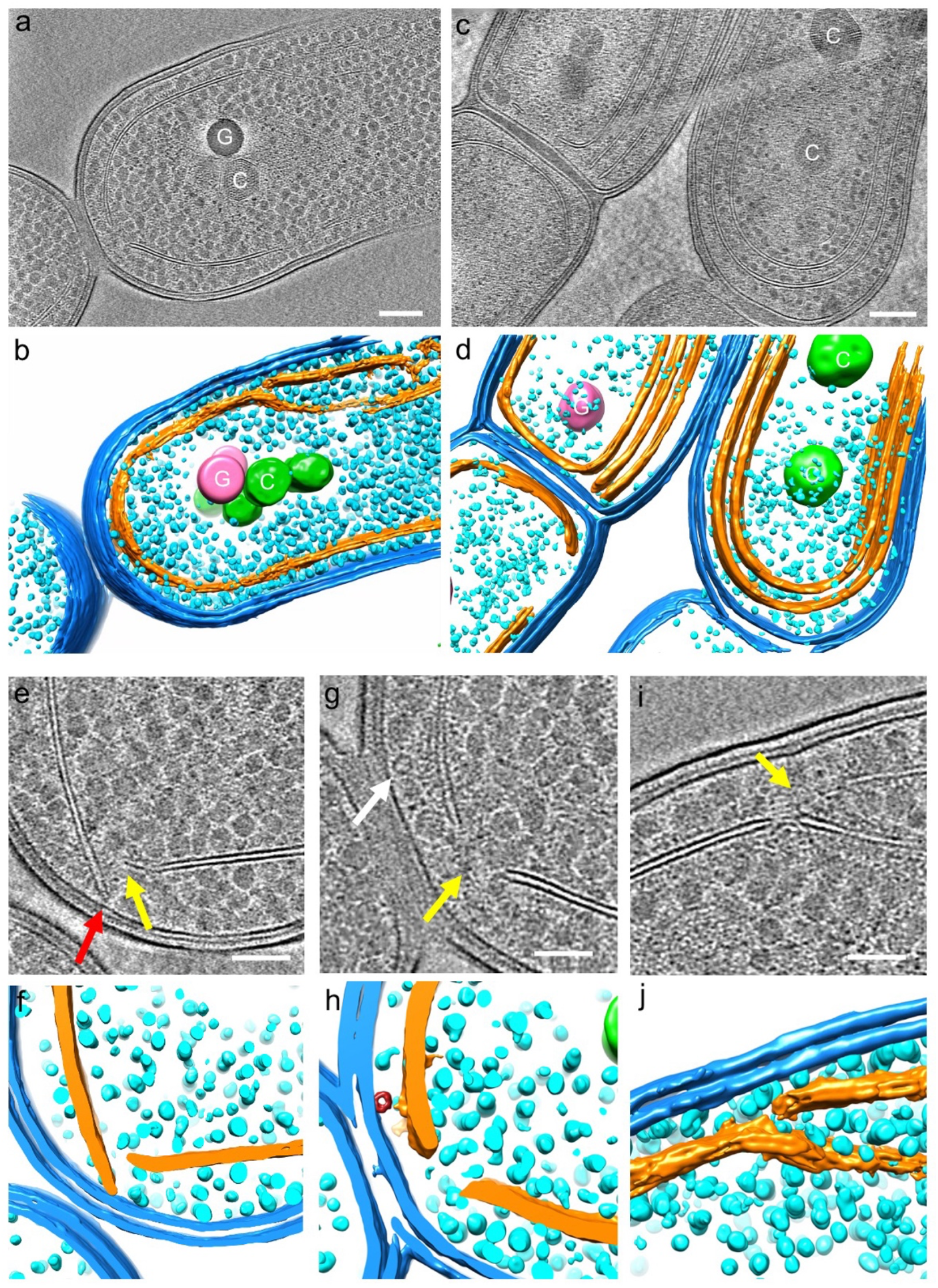
Cryo-electron tomography (ET) analysis of frozen-hydrated *Synechococcus* cell lamella. **a-d**, Tomographic slices (**a**,**c**) and segmented volumes (**b**,**d**) of cyanobacteria cells grown under high light (HL) (**a**,**b**) and these HL-grown cells transferred to low light (LL) condition for 1-2 days (**c**,**d**). **e-j**, Tomographic slices at high magnification (**e, g, i**) and segmented volumes (**f, h, j**) of cyanobacteria cells grown under HL. Outer and plasma membranes are in blue, thylakoid membranes in gold, glycogen granules in cyan, carboxysomes (“C”) in green, and large dense granules (“G”) in pink, most likely the polyphosphate bodies. Yellow arrows point to the gaps between two thylakoid membrane edges for the emergence of newly synthesized thylakoid membrane rafts (Supplementary Fig. 5). The red arrow points to the thylakoid tip located in the vicinity of the plasma membrane, but not connecting to it. The white arrow points to tubular structures. Scale bars, 200 nm in **a** and **c**, 100 nm in **e, g** and **i**. Single slice images have a thickness of 2.49 nm and the tomographic slices images are constructed from superimposed serial slices with the same thickness. Representative Cryo-ET images were derived from at least three biologically independent preparations with similar results.

Some small, tubular-like structures were also discerned close to the plasma membrane, with a diameter of 20-30 nm and a length varying between 30 and 48 nm (Fig. 2g, white arrow; Fig. 2h, red; Supplementary Fig. 6, 7; Supplementary Movie 1, 6). Some of these structures sit between the thylakoid tips and the plasma membrane and no apparent connection between these structures and the thylakoid membrane was determined (Fig. 2h, Supplementary Fig. 7, Supplementary Movie 1, 5). These unknown structures are reminiscent of the VIPP1 (Vesicle-Inducing Protein in Plastids 1, or Inner Membrane-associated protein of 30 kDa - IM30) multimers^22–27^ that are deduced to be related to thylakoid membrane biogenesis in cyanobacteria^28^ and chloroplasts^29^. Whether these tubular-like structures are Vipp1 assemblies or involved in thylakoid biogenesis remains to be explored.

Closer views of the tomogram revealed that the extra thylakoid fragment close to the plasma membrane did not form connections with the adjacent continuous thylakoid layer (also refers to the preexisting thylakoids) that is curved towards the central cytoplasm (Fig. 2i, 2j). This indicates that thylakoid biogenesis in *Synechococcus* occurs most likely at the peripheral sites of the cytoplasm close to the plasma membrane. Expansion of the new thylakoid layer likely pushes the existing thylakoid lamellae toward the central cytoplasm to develop sacs of thylakoid layers. The thylakoid membranes of *Synechococcus* are often discontinuous, as several perforations were observed in the thylakoid lamellae (Fig. 2g, 2h, Supplementary Fig. 8, Supplementary Movie 7), consistent with the previous electron tomography of thylakoids in *Synechococcus* and *Chlamydomonas*^17,30^. This membrane organization may provide a means for transport of ribosomes and synthesized proteins to specific cellular regions^16^. By contrast, the thylakoid perforations were not defined in the recent cryo-ET study of *Synechocystis*^15^.

### Sequential membrane integration of photosynthetic complexes during thylakoid biogenesis

To address how different photosynthetic complexes are integrated and assembled into thylakoid membranes during the redevelopment of thylakoids, we isolated the thylakoid membranes from the *Synechococcus* cells grown at different time points during thylakoid biogenesis. Label-free mass spectrometry^11,31^ was used to determine the composition and relative abundance of photosynthetic proteins integrated into the thylakoid membranes. Using mass spectrometry, it is possible to examine a vast number of target proteins, but the quality of obtained spectra may vary between proteins, especially the membrane-integral protein complexes. Thylakoid samples were digested and then loaded in mass spectrometry with an equal protein amount. A total of 1200 proteins were detected (Supplementary Data 1). The trajectories of the changes in protein abundance, normalized by the GL samples, were fitted well with fourth-degree polynomial functions (Fig. 3, Supplementary Table 1), likely due to the combination of protein biosynthesis and degradation processes.

**Figure 3.**
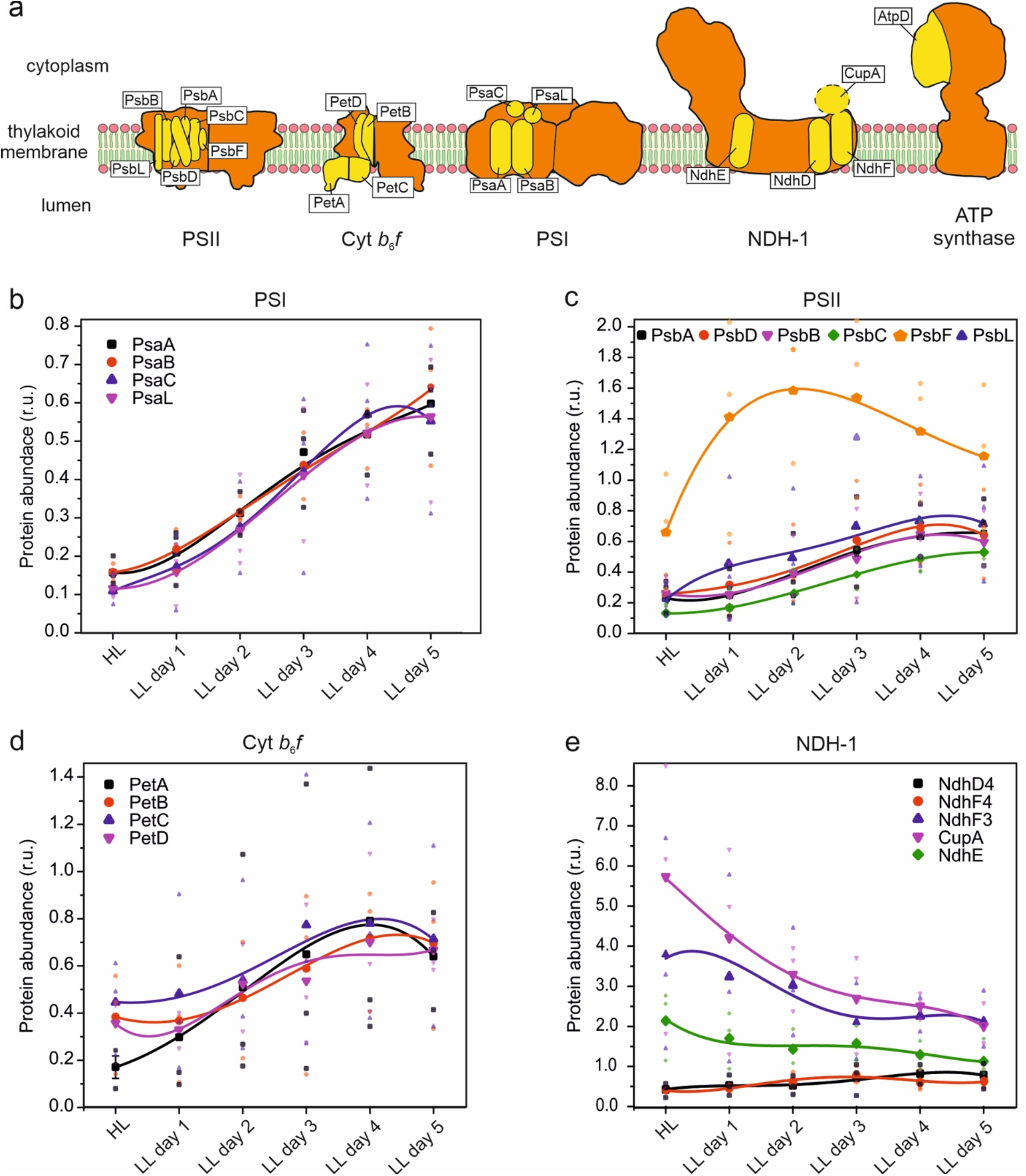
Expression of protein complexes residing in thylakoid membranes during thylakoid biogenesis. Membrane protein fractions were isolated from *Synechococcus* cells grown under GL, HL, and HL-grown cells transferred to LL for 5 days. **a**, Scheme of studied subunits from photosystem II (PSII), photosystem I (PSI), cytochrome *b*_6_*f* (Cyt *b*_6_*f*), type 1 NAD(P)H dehydrogenase-like complex (NDH-1) and ATPase. Illustration credit: Dr Tuomas Huokko, University of Liverpool; **b-e**, Relative protein quantification using label-free mass spectrometry data-dependent acquisition of subunits belonging to **b**, PSI; **c**, PSII; **d**, Cyt *b*_6_*f* and **e**, NDH-1. r.u., relative unit. Values are represented as means from 3 biologically independent experiments. All values are normalized to GL-samples. Transparent symbols represent three independent biological replicates. Fitted curves are fourth-degree polynomial functions. The corresponding adjusted R^2^ values for fitting are: PsaA, 0.98151; PsaB, 0.99498; PsaC, 0.99935; PsaL, 0.99967; PsbA, 0.99936; PsbD, 0.97432; PsbB, 0.96176; PsbC, 0.99982; PsbF, 0.99486; PsbL, 0.97164; PetA, 0.99612; PetB, 0.99497; PetC, 0.88053; PetD, 0.91018; NdhD4, 0.94908; NdhF4, 0.88885; NdhF3, 0.52203; CupA, 0.97552; NdhE, 0.61526. The equations of fitted curves are shown in Supplementary Table 1.

Proteomic results confirmed the reduced abundance of photosynthetic complexes in the HL-adapted thylakoid membranes (Fig. 3b-3d, Supplementary Data 1). The decrease in the PSI amount was represented by the reduced content of PSI reaction center proteins PsaA and PsaB, as well as PsaC (the docking site for ferredoxin or flavodoxin at the cytoplasmic side of PSI^32,33^) and PsaL (required for the formation of PSI trimer^34^) (Fig. 3b). When cells were transferred to LL, the relative content of PsaA, PsaB, PsaC and PsaL increased at comparable rates. After two days of LL adaptation, the PsaC and PsaL contents appeared to increase faster than those of PsaA and PsaB, corroborating the stepwise assembly of PSI proteins in PSI complex biogenesis^10^. Compared to other PSII subunits, the content of PsbF that forms part of cytochrome *b*_559_ was less reduced under HL, and there was a remarkable increase in the PsbF content at the initial stage of thylakoid biogenesis (Fig. 3c). A greater accumulation of PsbL also occurred at the initial thylakoid biogenesis process compared to PSII reaction center proteins PsbA (D1) and PsbD (D2), as well as the core antenna proteins PsbB (CP47) and PsbC (CP43). Thus, our results indicate that the stepwise biogenesis of PSII is promoted by the accumulation of PsbF and PsbL, which do not interact in the PSII holoenzyme, prior to D1 and D2^9^. Consistently, it has been shown that cytochrome *b*_559_ functions as a nucleation factor for the early steps of PSII assembly^35^ and the expression of the *psbEFLJ* operon is a prerequisite for D1 and D2 accumulation^36,37^. Among the Cyt *b*_6_*f* subunits, PetA (Cyt *f*) had a greater accumulation at the early step of thylakoid biogenesis compared to others (Fig. 3d), suggesting that cyanobacterial Cyt *b*_6_*f* also possesses a stepwise assembly.

Unlike the decrease in the abundance of photosynthetic complexes, NDH-1 complexes, which have different forms and participate in respiration, cyclic electron transfer and CO_2_ uptake in cyanobacteria^38^, were highly accumulated in thylakoid membranes induced by HL (Fig. 3e), consistent with previous studies^8^. NdhE and NdhV are common subunits for all the forms of NDH-1^39^, and their abundance under HL was noticeably higher compared to GL (Fig. 3e, Supplementary Fig. 9). Likewise, NdhF3 and CupA, which are the specific subunits of the low-CO_2_-inducible, high-affinity NDH-1_3_ complex for CO_2_ uptake^40^, were accumulated to levels 3.8 and 5.7 times than seen under GL, respectively (Fig. 3e). During thylakoid development, the relative amount of NdhE, NdhF3, CupA and NdhV gradually declined (Fig. 3, Supplementary Fig. 9). By contrast, the expression of NdhF4 and NdhD4, which are the specific subunits of constitutively expressed, low-affinity CO_2_ uptake system NDH-1_4_ complex^40^, was downregulated by ~60 % compared to GL (Fig. 3e), in agreement with previous transcriptional studies^41^; their relative amounts increased constantly up to the levels in the GL cells during thylakoid recovery (Fig. 3e). Our results demonstrate the distinct regulatory changes in the NDH-1 abundance during thylakoid biogenesis, which are dependent on the forms and functions of NDH-1.

The changes in the abundance of photosynthetic proteins during thylakoid biogenesis were further confirmed by sodium dodecyl sulphate-polyacrylamide gel electrophoresis (SDS-PAGE) combined with immunoblot analysis (Supplementary Fig. 9). Determination of ATPase composition and stoichiometry by mass spectroscopy was unreliable, due to high signal variation between biological samples. However, the specificity of the antibody against ATP-β, encoded by *atpD*, allowed us to monitor the changes in the ATPase content. The content of ATP-β subunits increased by 14% under HL compared to GL and was then gradually increased during LL adaptation (Supplementary Fig. 9), likely to meet the increasing demand of energy for thylakoid membrane biosynthesis.

To examine the assembly of functional photosynthetic complexes during thylakoid biogenesis, blue native-polyacrylamide gel electrophoresis (BN-PAGE) combined with immunoblot analysis was applied on the isolated thylakoid membranes. The contents of PSI trimers, PSII dimers and PSI/PSII monomers were decreased drastically under HL compared to GL, and then were increased during thylakoid biogenesis (Fig. 4a). During this process, PSI monomers accumulated faster than PSI trimers, PSII dimers and PSII monomers (Fig. 4b, 4c, 4d). Almost no PSI trimers were detected on LL day 1 (Fig. 4b, 4d), consistent with the lower abundance of PsaL that mediates PSI trimerization^34^ on LL day 1 (Fig. 3b). Along with thylakoid biogenesis, the relative amount of PSI trimers increased rapidly (Fig. 4b, 4d). Consistently, the monomer/oligomer ratio of PSI was higher than that of PSII at the early stage of thylakoid biogenesis and then became comparable at the later phase (Fig. 4e). These results indicated that during thylakoid biogenesis the assembly of PSI monomers in thylakoid membranes occurred before accumulation of PSI trimers and PSII complexes. PSI monomers were also detected in HL-adapted thylakoid membranes (Fig. 4b), ensuring PSI-mediated cyclic electron flow throughout cell growth^19^. In contrast, there were only a low amount of PSII monomers in the HL-adapted thylakoid membranes (Fig. 4c, 4d). Instead, the thylakoids accumulated a large amount of PSII core complexes lacking CP43, termed RC47 complexes^36,42^. On LL day 1, the RC47 complexes remained dominant in the PSII assemblies (Fig. 4c, 4d). During thylakoid biogenesis, PSII monomers and dimers accumulated at the expense of RC47, indicative of the stepwise assembly of PSII and the stoichiometric changes of distinct PSII assemblies. Recent studies showed that binding of the assembly factor Psb28 to RC47 protects PSII from photodamage during PSII assembly^43^.

**Figure 4.**
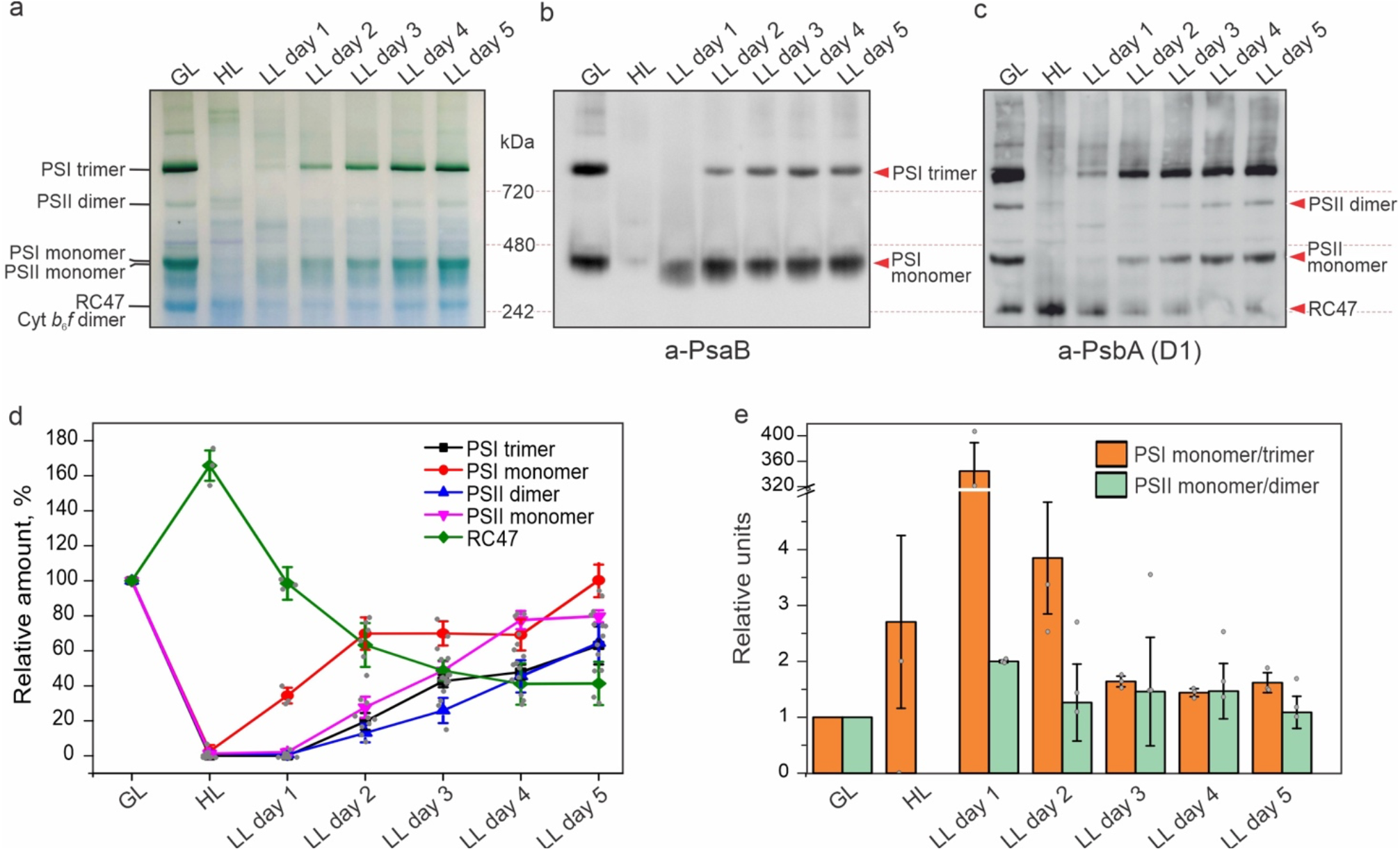
Oligomerization state of PSI and PSII during thylakoid membrane biogenesis in *Synechococcus*. **a**, BN-PAGE gel from isolated thylakoid membranes of cells grown under GL, HL, and HL-grown cells transferred to LL for 5 days. 200 µg of solubilized membrane fraction was loaded per lane. RC47: PSII core complex lacking CP43. **b**, BN-PAGE immunoblotted with antibody against PsaB. 50 µg of solubilized membrane fraction was loaded per lane. **c**, BN-PAGE immunoblotted with antibody against PsbA (D1). 50 µg of solubilized membrane fraction was loaded per lane. **d**, Relative quantification of protein amounts from BN-PAGE western blots. Values are means ± SD; *n* = 3 biologically independent experiments. **e**, Relative ratios of oligomeric and monomeric state of PSI and PSII during thylakoid membrane biogenesis. Values are means ± SD; *n* = 3 biologically independent experiments. Gel pictures and immunoblots are representative of 3 biologically independent experiments.

In agreement with the relative quantification of individual proteins (Fig. 3c) and levels of complexes (Fig. 4c), PSII activity was reduced almost 88% by HL treatment (Supplementary Fig. 10a) when most of PSII complexes are in the form of RC47 (Fig. 4c), in relative to GL. Under LL treatment, PSII activity gradually recovered when PSII monomers and dimers started to accumulate and become dominant (Supplementary Fig. 10a, Fig. 4c). After 4-day LL treatment, PSII activity reached about 87% of that in the GL-adapted cells. These results indicate that the recovery of PSII activity lags behind Chl synthesis (Supplementary Fig. l) and there might be a pool of Chls, which do not associate with functional photosystems in the recovering cells but could associate with IsiA forming dynamically the PSI-IsiA supercomplexes^11^ or could be redistributed between photosystems^44^. In contrast, the PSI activities were reported to be less variable than the PSII activity during the light-triggered thylakoid biogenesis^19^, consistent with our results indicating that the integration of PSI monomers into thylakoid membranes occurred before the assembly of PSII complexes during thylakoid biosynthesis (Fig. 3). Moreover, 77K fluorescence emission spectra with Chl excitation (435 nm) showed that the florescence at 695 nm specific for PSII was not distinguishable in the HL-grown cells but clearly reappears as a distinct peak in LL day 1 (Supplementary Fig. 10b). This indicates the biosynthesis of PSII and thylakoid membranes, as confirmed by the proteomics (Fig. 3c) and BN-PAGE results (Fig. 4). There was a blue-shift of the PSI fluorescence peak after HL phase (Supplementary Fig. 10b), consistent with the presence of a large amount of PSI monomers^45^ as observed in BN-PAGE (Fig. 4b). 77K fluorescence emission spectra, when excited at 600 nm, revealed that there was a notable increase in the fluorescence at 685 nm (phycobilisomes and PSII) relative to the fluorescence at 695 nm (PSII) under HL compared to GL (Supplementary Fig. 10c), indicating the functional decoupling of phycobilisomes and photosystems. A further increase in the phycobilisome fluorescence at LL day 1 suggested the presence of newly synthesized free phycobilisomes in cells. These phycobilisomes started to be energetically coupled with photosystems at LL day 2, when more functional PSII complexes were assembled in thylakoid membranes.

### Membrane location and redistribution of photosynthetic complexes during thylakoid biogenesis

We further utilized super-resolution confocal fluorescence imaging to visualize the *in situ* distribution of photosynthetic complexes during thylakoid biogenesis, using the fluorescently labeled *Synechococcus* mutants PSI-eGFP (tagged to PsaE), PSII-eGFP (tagged to PsbB (CP47)), Cyt *b*_6_*f*-eGFP (tagged to PetA), and ATPase-eGFP (tagged to AtpB)^12^. Cells at different thylakoid biogenesis stages were imaged by a Dragonfly spinning disk confocal microscope with the super-resolution radial fluctuations (SRRF)-stream technology^46^. The eGFP-tagged *Synechococcus* cells exhibited similar Chl/OD_750_ ratios as the WT cells under the corresponding light conditions (Supplementary Fig. 11), confirming that fluorescent tagging had limited effects on cell physiology as reported previously^12^. Under HL, PSI-eGFP exhibited both the thylakoid and cytosolic locations (Fig. 5a), probably due to the HL-induced detachment of the peripheral PSI subunit PsaE from thylakoids^47^ that was tagged with eGFP^12^. This was confirmed by immunoblot analysis (Supplementary Fig. 12). Transferring the HL-adapted cells to LL rescued the thylakoid location of PSI-eGFP (Fig. 5a, Supplementary Fig. 12). At the initial stage of thylakoid biogenesis, the Chl fluorescence in all studied strains was brighter on one longitudinal side of the cell than the other side (Fig. 5), demonstrating the asymmetrical biosynthesis of thylakoid membranes as revealed in thin-section EM (Fig. 1a, 1b) and cryo-ET (Fig. 2). The PSI, PSII, Cyt *b*_6_*f*, and ATPase also displayed the asymmetrical distribution on LL day 1 and GFP signals were concentrated on the same longitudinal side of the cells as Chl signals (Fig. 5, Supplementary Fig. 13), further confirming the asymmetric biosynthesis of thylakoid membranes in *Synechococcus* cells. Compared to the other three photosynthetic complexes at this stage, PSII showed a higher degree of asymmetrical distribution (Fig. 5c, white arrows; Fig. 5d, Supplementary Fig. 13), suggesting that a great amount of PSII complexes were integrated into the newly synthesized thylakoid membranes. During the synthesis of thylakoid membranes, PSI displayed more clustered distribution in mature thylakoid membranes, frequently at the poles of thylakoids (Fig. 5a, orange arrows; Fig. 5b), giving rise to the more uneven distribution in thylakoid membranes (Supplementary Fig. 14a), whereas PSII, Cyt *b*_6_*f*, and ATPase complexes exhibited relatively even distribution in thylakoid membranes at the later stage of thylakoid biogenesis (Fig. 5, Supplementary Fig. 14b-14d). These results provide evidence for the membrane redistribution of photosynthetic complexes during thylakoid biogenesis, ensuring formation of specific membrane domains for function^1,8,12^. Likewise, phycobilisomes colocalized largely with Chl fluorescence during thylakoid membrane reconstruction (Supplementary Fig. 15).

**Figure 5.**
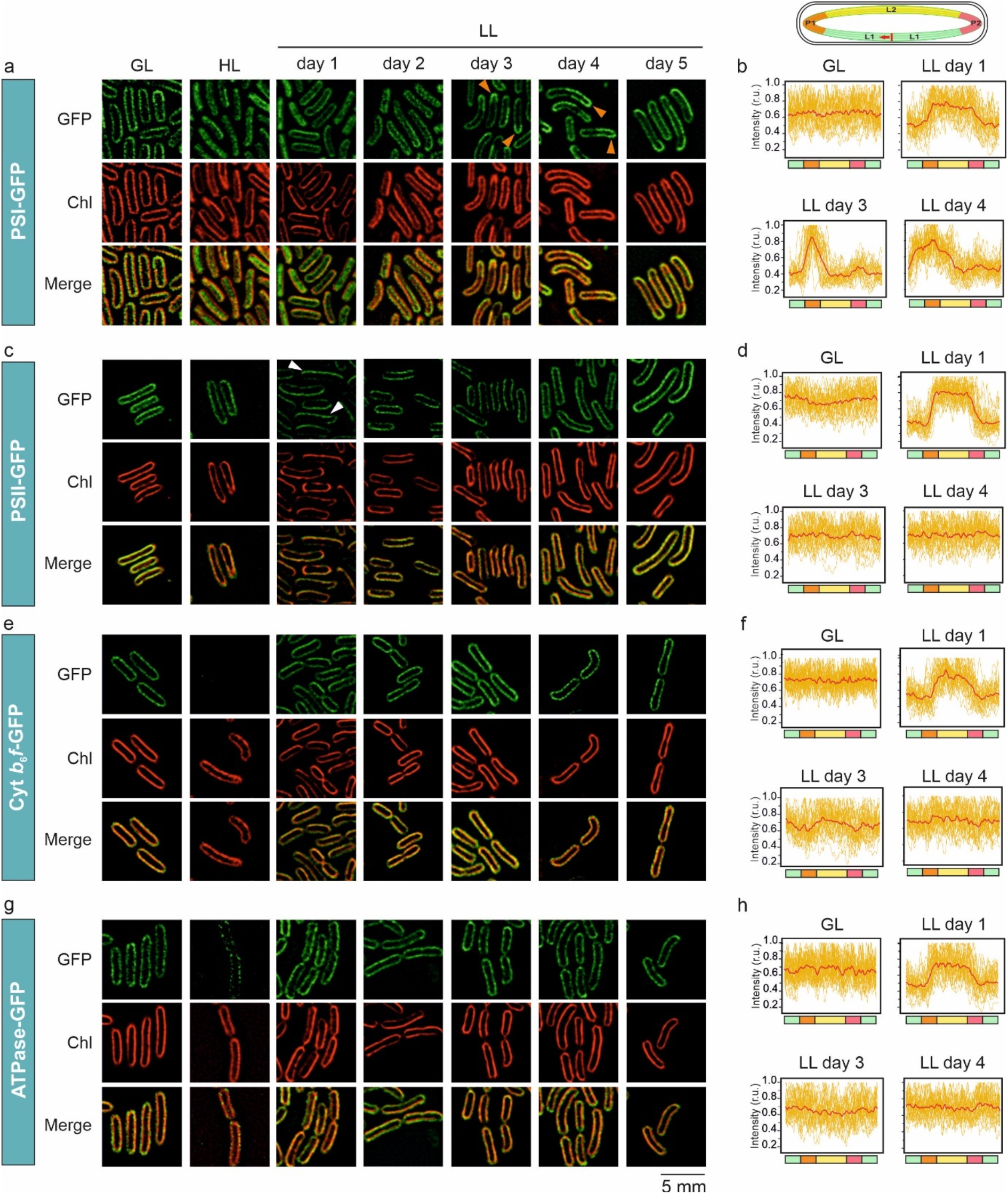
Expression and intracellular localization of photosynthetic complexes *in vivo* in *Synechococcus* during thylakoid membrane biogenesis. **a**, PSI:eGFP; **c**, PSII:eGFP; **e**, Cyt *b*_6_*f*:eGFP and **g**, ATPase:eGFP *Synechococcus* cells imaged by Dragonfly spinning disk confocal microscope with the super-resolution radial fluctuations (SRRF)-stream technology. Figures are representative of 3 biologically independent experiments. First row: GFP emission, second row: Chl fluorescence, third row: merge. **b, d, f**, and **h**, GFP profile analysis of PSI-, PSII-, Cyt *b*_6_*f*- and ATPase-eGFP around the cell in these strains. Profiles in red are averages from 30 individual imaged cells, shown in yellow on the background, from 3 biologically independent experiments. In the schematic model shown on the top, L1 in green: longitudinal cell side 1, P1 in orange: pole 1, L2 in yellow longitudinal cell side 2, P2 in pink: pole 2. In each cell tracking of eGFP-signal was started from the same site (indicated by red vertical line) and to the same direction (indicated by red arrow). A higher difference between minimum and maximum of signal indicates more clustered distribution of proteins. Cells were grown under growth light (GL), high light (HL) and HL-grown cells were then transferred to low light (LL) conditions for five consecutive days.

## Discussion

In this study, we systematically characterized the dynamic process of thylakoid membrane biogenesis in *Synechococcus*. By applying classical chemical fixation and thin-section EM of cyanobacterial cells^9,48–52^, we showed the regulation of thylakoid membrane abundance in cyanobacteria in response to the changing light intensity during cell growth. Consistent with the finding derived from conventional EM, our study by combining cryo-focused ion beam (cryo-FIB) milling and in situ cryo-ET further revealed that no physical fusion was discerned between the plasma membrane and regular thylakoid concentric cylinders *in vivo*, in line with recent observations on *Synechocystis*^15^. The newly synthesized thylakoid membrane layers appear asymmetrically on one longitudinal side of the cell between the plasma membrane and pre-existing thylakoids. The assembly and integration of photosynthetic complexes in thylakoids proceed via a stepwise fashion and vary in temporal order and spatial location. The integrated photosynthetic complexes are reorganized in membranes in the course of thylakoid biogenesis. These results allow us to propose a model to illustrate the biogenesis pathway of thylakoid membranes in *Synechococcus* (Fig. 6).

**Figure 6.**
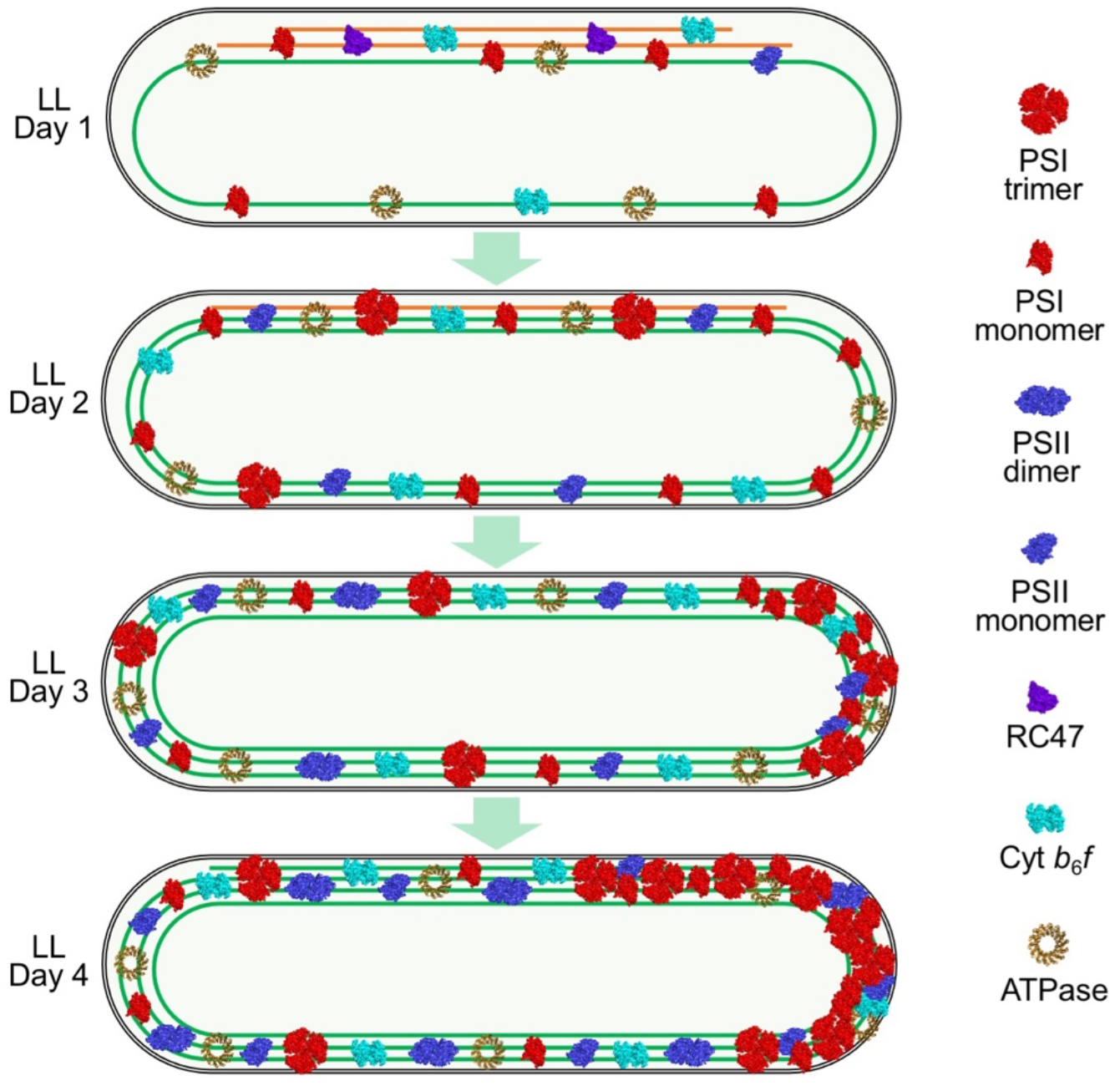
A schematic model representing the generation and distribution of both thylakoid membrane and photosynthetic protein complexes, including their oligomerization state, at different stages of thylakoid biogenesis in *Synechococcus*. Note that the number of individual complexes has no direct links to their real stoichiometry *in situ*. Existing membrane parts are in green and new ones in orange during the initial steps of thylakoid biogenesis process. The schematic models of photosynthetic complexes were generated based on their PDB structures. PDB ID: PSI trimer and monomer, 1JB0; PSI dimer and monomer and RC47, 3WU2; Cyt *b*_6_*f*, 2E74; ATPase, 6FKF.

Where thylakoid biogenesis initiates and how the biogenesis process proceeds are poorly understood. Three distinct models for cyanobacterial thylakoid membrane biogenesis are currently being debated^53^: (1) The synthesis of photosynthetic components occurs at the plasma membrane before migrating to thylakoids via their fusion sites, based on membrane fractionation combined with immunoblot and proteomic studies^54,55^; (2) The synthesis of photosynthetic components (including protein, pigment, and lipid) occurs *in situ* in pre-existing thylakoid membranes; (3) The photosynthetic components are synthesized at ribosome-rich sites and are assembled in specialized regions in the thylakoid membrane^14^. The thylakoid membrane structures in cyanobacteria vary among different species. It has been well documented that in the spherical unicellular cyanobacterium *Synechocystis*, thylakoid membranes converge at several sites close to the plasma membrane^14,17^. These convergence regions may serve as the biogenic sites of thylakoids^56^. It has remained unclear how thylakoid biogenesis is initiated in rod-shaped *Synechoccoccus* cells that have regular thylakoid sacs that form concentric cylinders without convergence sites (Fig. 1, 2).

Our cryo-ET results showed no direct connections between the plasma membrane and regular thylakoid membranes in *Synechococcus* at the initial stage of thylakoid biogenesis (Fig. 2, Supplementary Figs. 3 and 4), similar to the observations in *Synechocystis*^15^. Moreover, we observed the extension of thylakoid lamellae towards the plasma membrane, and the thylakoid tips are in close vicinity of the plasma membrane (Fig. 2, Supplementary Fig. 5, Supplementary Movie 3), reminiscent of the thylapses characterized in *Synechocystis*^15^. To our knowledge, this is the first characterization of the “thylapse” - like structures in the cyanobacterial cells possessing regular thylakoid cylinders. Whether these “thylapse” - like structures are involved in thylakoid biogenesis remains to be experimentally verified. It is worth noting that the thylapse structures observed in *Synechoccoccus* are mainly located at the cell poles, and thus are not likely to be involved in cell division.

We also observed that additional thylakoid layers representing the newly synthesized thylakoids appear specifically on one longitudinal side of the cell, resulting in the asymmetrical distribution of thylakoid lamellae at the early stage of thylakoid biogenesis (Fig. 1, 2, 5). These thylakoid sheets sit between the plasma membrane and the pre-established continuous thylakoid layer (Fig. 1, 2), suggesting that thylakoid biosynthesis in *Synechoccoccus* likely commences from the regions close to the plasma membranes rather than from the inner cytoplasmic side of the cell. Consistent with this, some small membrane segments that have one side connected with the thylakoid lamellae and the other end approaching the plasma membrane have been seen (Supplementary Fig. 5, Supplementary Movies 4 and 5). Likewise, some tube-like structures were observed next to the plasma membrane and sometimes close to the thylakoid tips (Fig. 2g, 2h, Supplementary Figs. 6 and 7, Supplementary Movies 1 and 6). The extension of the peripheral thylakoid fragments will compress the cytoplasmic space (Fig. 1a, 2b, 2j), resulting in the development of thylakoid sacs or the spiral organization towards the central cytoplasm. Therefore, a more symmetrical, regular concentric arrangement of thylakoid membranes in *Synechoccoccus* was established at the later stage of thylakoid biogenesis (Fig. 1, 5, 6).

Identification and relative quantification of protein complexes in isolated thylakoid membranes during the development of thylakoid membranes revealed the sequences of the assembly and membrane insertion of photosynthetic complexes into thylakoids (Fig. 3, Supplemental Fig. 9). The biosynthesis of thylakoids starts from the reduced thylakoid membranes acclimated in HL, in which there are relatively low amounts of PSI and PSII (Fig. 3b and 3c), a reduced Cyt *b*_6_*f* content (Fig. 3d), a relatively comparable amount of ATPase (Supplementary Fig. 9), as well as an increased level of NDH-1_3_ and a lower amount of NDH-1_4_ (Fig. 3e), compared to those in GL. Under HL, PSI complexes are mainly in the monomeric form whereas most of PSII are present as RC47 complexes (Fig. 4b, 4c) with a low PSII activity which should be derived from a low level of PSII monomers (Fig. 4, Supplementary Fig. 10). The PSI/PSII ratio in *Synechococcus* is 4.5^12^. When thylakoid biogenesis initiates, the assembly of PSI complexes occurred earlier than that of PSII (Fig. 4). Moreover, PSI monomers undertook a faster accumulation than PSI trimers and PSII holoenzymes. Recent AFM images have demonstrated that PSI trimers, monomers, and some dimers coexist in the *Synechococcus* thylakoid membranes^11^. The ratio of PSI monomer/trimer varied during thylakoid biogenesis, along with the changes in the PsaL content (Fig. 3b, 4b, 4d, 4e). These results outlined the stepwise *de novo* assembly process of PSI and the flexibility of PSI oligomerization that is regulated by PsaL^34^. The functional discrepancy of different PSI oligomers and how their stoichiometries are physiologically regulated in cyanobacterial cells remain elusive. AFM also revealed the association of PSI with PSII in native thylakoid membranes^11^, highlighting the possible roles of PSI in PSII biogenesis^57^ and photoprotection via efficient energy spillover^58^.

The temporal stepwise assembly of photosynthetic complexes from building modules was also demonstrated by the early accumulation of PsaA and PsaB compared to other PSI components (Fig. 3b), and there was a notable increase in the relative amounts of PsbF and PsbL compared to the PSII core proteins D1 and D2 during the initial stages of thylakoid biogenesis (Fig. 3c). As D1 protein is strongly involved in PSII photoinhibition, the lack of a considerable increase in the D1 amount at the early stage of thylakoid biogenesis could reflect the balanced synthesis and degradation of D1 in the PSII repair cycle^59,60^. Moreover, a large amount of RC47 complexes were accumulated under HL and LL day 1, prior to the formation of PSII monomers and subsequent dimers that are considered as the final photochemically active form of PSII^9,59^ (Fig. 4c, 4d).

NDH-1 complexes in cyanobacterial thylakoid membranes are crucial for both photosynthetic cyclic electron transfer and respiration^38^. *In vivo* observations illustrated that in thylakoid membranes, NDH-1 complexes could reorganize to be clustered in respiratory membrane zones under a lower light to facilitate respiratory electron flow, or to be distributed throughout the thylakoid membranes adjacent to PSI for cyclic electron transfer^8^. In addition to the biochemical evidence^61^, the tight association of NDH-1-PSI supercomplexes has been directly visualized in cyanobacterial thylakoid membranes^11^. Cyanobacteria possess different isoforms of NDH-1 that vary in protein composition and function^62^. Our results indicated that the stoichiometric regulation of different NDH-1 isoforms in response to changes in light conditions differs. The total NDH-1 complexes, including the low-CO_2_-inducible, high-affinity NDH-1_3_, were accumulated under HL, in agreement with the previous studies^8,11^, and then the relative amount decreased during the LL-induced thylakoid biosynthesis (Fig. 3e). In contrast, the constitutively expressed, low-affinity NDH-1_4_ exhibited sharply opposite responses to light regulation. The underlying mechanisms and physiological significance in CO_2_ assimilation remain to be determined.

In addition to the assembly orders of photosynthetic complexes, our study also provides clues about the *de novo* membrane integration and assembly sites of photosynthetic complexes and the dynamics of their subcellular locations. The thylakoid membranes^63^, thylapse membrane domains^15^ or PratA-defined membrane (PDM) regions^56,64–66^ in *Synechocystis*, and the translation zones in the chloroplast of *Chlamydomonas*^67,68^ have been proposed to be the principal sites for the initial biogenesis and/or assembly of photosynthetic complexes, in particular PSII. Our confocal images showed explicitly that at the initial step of thylakoid biogenesis, compared to other photosynthetic complexes, PSII complexes exhibited a greater accumulation on one longitudinal side of the *Synechoccoccus* cell where the newly synthesized thylakoid fragments appear (Fig. 5, Supplementary Figs. 13 and 14). Since eGFP was tagged to CP47^12^ and the PSII complexes appeared predominantly as RC47 at the initial stage of thylakoid biogenesis (Fig. 4c, 4d), our results indicate that the initial steps of RC47 assembly occur predominantly in these newly generated thylakoid fragments. In a very recent study, we have revealed that the mRNA molecules encoding the core subunits of PSI and PSII clustered at the thylakoid surfaces that are mainly adjacent to the central cytoplasm^66^. We showed that the locations of mRNA molecules correspond to the sites of translation. After translation, these synthesized photosystem proteins could be directed, through unknown mechanisms, to the newly formed thylakoid layers at the peripheral side of the cytoplasm. Indeed, the perforations in the thylakoid lamellae that we observed in cryo-ET (Fig. 2g, 2h, Supplementary Fig. 8, Supplementary Movie 7) could facilitate rapid protein transport between spatially distinct cellular regions through the thylakoid membranes. During the expansion of thylakoid membranes, PSII assemblies spread throughout thylakoid membranes (Fig. 5c, 5d, Supplementary Fig. 14) and heterogeneous clustering and arrays of PSII dimer rows have been observed in the thylakoid membranes^11,12^. Consistently, FtsH complexes involved in D1 degradation exhibit a patchy distribution along the thylakoid membranes of *Synechocystis*, suggesting the presence of PSII repair “zones” ^69^.

Unlike PSII, the initial membrane insertion and assembly of PSI complexes seem to occur more evenly in both newly synthesized and existing thylakoid membranes (Fig. 5a, 5b, Supplementary Fig. 13), suggesting that there might be different protein targeting mechanisms or assembly sites of PSI and PSII. During the thylakoid biogenesis process, PSI complexes could redistribute to exhibit more polar membrane distribution (Fig. 5a, 5b, Supplementary Fig. 14a), consistent with the previous studies^12^. The PSI-enriched thylakoid membranes of *Synechococcus* have been observed in previous AFM studies^11^. Therefore, our results provide evidence for the dynamic membrane location of photosynthetic and respiratory complexes during thylakoid biogenesis.

In summary, our study provides insights into the spatial-temporal assembly and biogenesis of photosynthetic complexes and thylakoid membranes in the rod-shaped unicellular cyanobacterium *Synechococcus*. We demonstrate that thylakoid membrane formation is a highly dynamic process, involving the assembly and functional modulation of photosynthetic protein complexes. It is still unclear how pigment and lipid biosynthesis are functionally coordinated with protein biosynthesis and assembly during thylakoid biogenesis. We believe that the methodologies are suited to assess the biogenesis and development of morphologically distinct thylakoid membranes in diverse cyanobacterial species and chloroplasts as well as other biological membrane systems.

## Methods

### Strains and culture conditions

*Synechococcus elongatus* PCC 7942 (WT) and eGFP-tagged *Synechococcus* strains^12^ were grown in BG11 medium under the continuous white light of 40 µmol photons m^−2^ s^−1^ at 30°C in culture flasks with constant shaking. Experimental cultures were grown in a photobioreactor Multi-Cultivator MC 1000 (Photon Systems Instruments, Brno, Czech Republic). Parallel 80-ml cultures were started from pre-experimental cultures by inoculating cells to the starting OD_750_ = 0.2 in fresh BG11 medium. Cells were first cultured at 30°C for 14 days either under 40 µmol photons m^−2^ s^−1^ (growth light, GL) or under 300 µmol photons m^−2^ s^−1^ (high light, HL). After this, OD_750_ of the HL culture was adjusted to 0.6 with fresh BG11 and transferred to under 20 µmol photons m^−2^ s^−1^ (LL, low light) for 5 days. Optical densities were measured using DS-11 FX+ spectrophotometer (DeNovix). For physiological and structural measurements, as well as for protein extraction, cells were harvested after 14 days of growth under both GL and HL conditions along with five consecutive days after transferring cells from HL to LL conditions.

### Chl extraction and determination by methanol extraction

Chl was extracted from cells with 90% methanol, and Chl concentrations were determined by measuring OD_665_ and multiplying it with extinction coefficient factor 12.7^70^.

### Thylakoid membrane extraction, protein electrophoresis, and immunoblot analysis

The total protein extracts of the *Synechococcus* cells were obtained according to previous methods^71,72^: cells suspended in isolation buffer (50 mM HEPES-NaOH, pH 7.5, 30 mM CaCl_2_, 800 mM sorbitol, and 1 mM ε-amino-n-caproic acid) were disrupted by vortexing with glass beads (212–300 μm in diameter) at 4°C and centrifuged at 3,000× g for 5 min to remove the glass beads and unbroken cells. From total protein extracts thylakoid membranes and soluble proteins were fractioned as described previous^12^: membranes were pelleted by centrifugation at 17,000× g for 30 min and resuspended in storage buffer (50 mM Tricine-NaOH, pH 7.5, 600 mM sucrose, 30 mM CaCl_2_, and 1 M glycinebetaine).

Protein complexes in their native form from isolated membrane fractions of *Synechococcus* were studied by Blue Native (BN)-PAGE according to previous methods^72^: isolated membranes were washed with washing buffer (330 mM sorbitol, 50 mM Bis-Tris, pH 7.0, and 250 µg mL^−1^ of Pefabloc, Sigma) and suspended in 20% glycerol (w/v), 25 mM Bis-Tris, pH 7.0, 10 mM MgCl_2_, and 0.01 unit mL^−1^ RNase-Free DNase RQ1 (Promega) at the final concentration of 20 µg protein µL^−1^. The samples were incubated on ice for 10 min, and the equal volume of 3% n-dodecyl-b-D-maltoside was added. Samples were first solubilized for 10 min on ice and after that for 20 min at room temperature following centrifugation at 18,000× g for 15 min to remove insoluble material. The supernatant was collected and mixed with 1/10 volume of 0.1 M EDTA and 1/10 volume of sample buffer (5% Serva blue G, 200 mM Bis-Tris, pH 7.0, 75% sucrose, and 1 M ε-amino-n-caproic acid). Samples were applied to NativePAGE Bis-Tris protein gels with 4-16% gradient (Thermo Scientific) and voltage was gradually increased from 50 V up to 200 V during the gel run. After electrophoresis, the proteins were transferred to a PVDF membrane (Immobilon-P; Millipore) and examined with protein-specific antibodies; α-PsaB (AS10 704 Agrisera) in dilution of 1:1000 and α-PsbD (AS06 146, Agrisera) in dilution of 1:4,000. Secondary antibody (AS09 602, Goat anti-Rabbit IgG (H&L), HRP conjugated, Agrisera) was applied in dilution of 1:10,000. 200 µg of proteins from the thylakoid membrane fraction per sample were loaded to BN-PAGE gels. For immunoblot analysis, 50 µg of proteins from the thylakoid membrane fraction per sample were loaded to BN-PAGE gels.

For denatured electrophoresis, crude thylakoid membrane proteins were denatured according to previous methods^73^; Sodium dodecyl sulfate (SDS) sample buffer (200 mM Tris-HCl, pH 6.8, 8% SDS, 400 mM dithiothreitol (DTT), 0.02% bromophenol blue) was added to thylakoid membrane samples and incubated at 75°C for 10 min. Then, proteins were separated by 15% (w/v) SDS-PAGE, transferred to a PVDF membrane (Immobilon-P, Millipore) and analyzed with the protein-specific antibodies; α-PsaB (AS10 695, Agrisera) in dilution of 1000, α-PsbA (AS05 084, Agrisera) in dilution of 1:4000, α-PsbD (AS06 146, Agrisera) in dilution of 1:4,000, α-PetC (AS08 330, Agrisera) in dilution of 1:1500, α-AtpD (AS05 085, Agrisera) in dilution of 1:3,000, α-NdhV (a generous gift from Dr. Hualing Mi) in dilution of 1:4000 and α-GFP (A-11122, Invitrogen) in dilution of 1:5,000. Secondary antibody (AS09 602, Goat anti-Rabbit IgG (H&L), HRP conjugated, Agrisera) for all other primary antibodies besides for α-GFP was applied in dilution of 1:10,000 and secondary antibody (AS11 1772; Goat anti-Mouse IgG, HRP Conjugate, Agrisera) for α-GFP in dilution of 1:10,000.

### Transmission electron microscopy

For transmission electron microscopy (EM) analysis samples were prefixed with 4% paraformaldehyde and 2.5% glutaraldehyde in 0.05 M sodium cacodylate buffer (pH 7.2). *Synechococcus* cells were then postfixed with 1% osmium tetroxide, dehydrated with increasing alcohol concentrations (30 to 100%), embedded in resin and cut to thin sections (70 nm)^48–51^. These were stained with 4% uranyl acetate and 3% lead citrate. Images were recorded using a Tecnai G2 Spirit BioTWIN transmission electron microscope (Field Electron and Ion Company, FEI) equipped with a Gatan Rio 16 camera. Distance between thylakoid membrane layers was determined from TEM-images with FIJI image processing package (ImageJ, NIH).

### Cryo-electron tomography

Cell vitrification and cryo-focused ion beam (cryo-FIB) milling: *Synechococcus* cells were diluted to OD_750_ = ~0.8 in their culture medium before plunge freezing. 3 µl of suspended cells in culture medium were applied to the front side of glow-discharged R2/2 carbon-coated copper grids (Quantifoil MicroTools), with additional 1 µL cells added to the back side. The grids were blotted for 4 seconds before vitrified in liquid ethane using a Leica GP2 plunge freezer. The vitrified grids were subsequently loaded to an Aquilos focused ion beam/scanning electron microscope (FIB/SEM) (Thermo Fisher Scientific) to prepare thin lamellae of cells. To reduce overall specimen charging, the grids were sputter-coated with platinum before milling. The ion beam current was gradually adjusted to lower values as the lamellae thinning progress (first using 0.1 nA until 0.75 µm thick, and then 50 pA until ~ 250 nm thick).

Cryo-electron tomography data collection: Tomography tilt-series were acquired from lamella using a Titan Krios microscope (Thermo Fisher Scientific) operated at 300 kV. The tilt-series data were acquired with a stage pre-tilt of −7º to account for the angle of lamella, making the lamella surface roughly perpendicular to the electron beam. Tilt-series were collected from −42º to 42º in 2º increment by a group of 3 using dose symmetric scheme^74^ in SerialEM^75^, with a target defocus of −10 µm. The tilt series reconstruction was performed with a binning factor of 4, resulting in a final pixel size of 2.492 nm. Micrographs were recorded with a K3 camera equipped with a Gatan Quantum energy filter operated in zero-loss mode with a 30 eV slit width. The exposure time was set to 5 seconds and movies were fractionated into 40 subframes. The nominal magnification of the recorded images was 15000×, with a pixel size of 6.23Å. The total accumulated dose was ~120 e^−^/Å^2^.

Tomogram reconstruction and segmentation: The movie frames were motion-corrected using MotionCor2 with 5 × 5 patches^76^. Tilt-series alignment was performed using IMOD^77^. The black dot-like stains on the surface of lamella were selected as fiducial markers, which were manually traced to create the fiducial model. Tomogram was reconstructed using 4× binned (24.92 Å/pixel) micrographs without CTF correction with 5 cycles of SIRT^78^. The automated segmentation/annotation of 3D volumes was performed using neuronal network implemented in EMAN2.3^79^, with additional manual adjustments. These volumes were visualized in 3D using UCSF Chimera^80^.

### Mass spectroscopy

In-solution digestion of thylakoid membrane samples for MS: Thylakoid membrane pellets were reconstituted, reduced, alkylated, and digested as described in^11^, with the exception that trypsin to protein ratio was 1:50. The following day Rapigest was removed by the addition of 0.5% (v/v) TFA and incubation at 37°C for 45 min. Digests were centrifuged at 17,200× g for 30 min and the clarified supernatants aspirated. Samples were stage– tipped on C18 filters to remove chlorophyll prior to LC-MS/MS analysis.

LC-MS/MS analysis: Data-dependent LC-MS/MS analyses were conducted on a QExactive quadrupole-Orbitrap mass spectrometer coupled to a Dionex Ultimate 3000 RSLC nano-liquid chromatograph (Dionex/Thermo Fisher Scientific). An equivalent of 100 ng peptides per sample were injected for mass spectrometry. The mass spectrometer was operated in DDA mode with survey scans between *m*/*z* 300-2000 acquired at a mass resolution of 70,000 (FWHM) at *m*/*z* 200. The 10 most intense precursor ions with charge states of between 2+ and 5+ were selected for MS/MS with an isolation window of 2 *m*/*z* units.

Database search and Protein identification: Raw data files were searched against the UniProt proteomes database of *Synechococcus elongatus* 7942 (UniProt ID: UP000002717) using Proteome Discoverer software (Thermo Fisher Scientific version 1.4.1.14) connected to an in-house Mascot server (Matrix Science, version 2.4.1). A precursor ion tolerance of 10 ppm and a fragment ion tolerance of 0.01Da were used with carbamidomethyl cysteine set as a fixed modification and oxidation of methionine as a variable modification.

Label-free quantification in Progenesis QI for MS: Raw mass spectral data files were processed using Progenesis-QI (v4.1; Nonlinear Dynamics) to determine total protein abundances. All raw files were initially automatically aligned, according to retention time, to produce an aggregate LC-MS map, from which peptide feature charge-states +1 and > +7 were excluded. A precursor ion tolerance of 10ppm and a fragment ion tolerance of 0.01Da were used, with carbamidomethylation of cysteine set as a fixed modification and oxidation of methionine as a variable modification. Trypsin was the specified enzyme, and one missed cleavage was allowed. A peak list was exported to Mascot and searched against the UniProt proteomes database of *Synechococcus elongatus* 7942 (UniProt ID: UP000002717) using the Mascot search engine (version 2.4.1; Matrix Science, UK).

### Confocal fluorescence microscopy and imaging analysis

To immobilize cells for imaging 10 µl of culture was pipetted on top of agar and dried drop was cut out and placed against cover slip. Live-cell super-resolution spinning disc confocal fluorescence imaging was performed on a Dragonfly microscope (Andor) utilizing super-resolution radial fluctuations (SRRF)-Stream technology. 63× oil-immersion objective (numerical aperture: 1.46) and excitation at 488 nm were used in imaging. Applied setting for SRRF was: SRRF frame count = 100, SRRF radiality magnification = 4, SRRF ring radius = 1.00 px, SRRF temporal analysis = mean, symmetrical binning 1×1. The confocal pinhole was set to 40 μm. GFP and chlorophyll signals were detected using a 488-nm laser, 525-nm filter (bandwith: 50 nm) and a 700-nm filter (bandwith: 75 nm), respectively. The phycobilisome fluorescence signal was detected using a 561-nm laser and a 620-nm filter (bandwith: 60 nm). Images were processed with FIJI image processing package.

To obtain the GFP-signal intensities around the cell, a line was traced following the cell outline and fluorescence was plotted as a function of normalized line distance. Signal tracing was always started in the middle of the longitudinal side of the cell. The average curve was generated by Origin software (OriginLab Corporation). The patchiness in the distribution of GFP-tagged photosynthetic protein complexes along thylakoid membranes we analyzed using the method described in^8^. A line was traced around the thylakoids and fluorescence was plotted as a function of distance along the line. The standard deviation of the GFP signal profile was then normalized by the total GFP fluorescence. Evenly distributed fluorescence demonstrates low standard deviation whereas patchy fluorescence results in higher standard deviation. To examine the distribution of GFP-tagged photosynthetic protein complexes between longitudinal cell sides at the early stage of thylakoid biogenesis, the maximum fluorescence from both cell sides was traced to obtain their ratio. Colocalization analysis of emission signals was conducted with Coloc2 in FIJI image processing package.

### Cell counting

Cells were counted on a hemocytometer slide using cell suspensions at about 1-5 μM Chl a depending on the stage of thylakoid regeneration.

### Whole-cell absorption spectra and inherent Chl determination

Whole-cell absorption spectra were measured at room temperature with 1 nm increments using a modernized Aminco DW2000 UV/Vis spectrophotometer (Olis, USA), which places the sample cuvettes very close a large detector window, hence minimizing distortions due to turbidity. Chlorophyll concentration was estimated from the cell absorption spectra using the formulae as described previously^81^: *A*Chl_678_ = 1.0162 × A_678_ - 0.0630 × A_625_ and Chl was estimated from A_678_ by an absorption coefficient of 68 mM^−1^ cm^−1^. The raw absorption values were below 0.5 at 750 nm and below 0.8 at the 620/680 nm peaks used to calculate pigment content. For presentation, absorption spectra were normalized to turbidity measured at 750 nm.

### 77K fluorescence emission spectra

Fluorescence emission spectra were recorded at 77K with a Perkin-Elmer LS55 luminescence spectrometer equipped with a liquid nitrogen housing. Cells were resuspended to 5 µM Chl in BG11 medium. Cell suspensions were then loaded into silica capillary tubes and dark-adapted for 5 min before freezing by plunging into liquid nitrogen. Spectra were recorded for the frozen samples with excitation at 435 nm (for Chl *a* excitation) or 600 nm (for phycocyanin excitation) and emission at 620-750 nm. The excitation slit-widths were 5 nm with 435 nm excitation and 3 nm with 600 nm excitation. Emission spectra were corrected for the instrument spectral response and normalized after subtracting the background signal. Spectra with 435 nm excitation were normalized to the PSI emission peak (~ 720 nm) and spectra with 600 nm excitation were normalized to the phycocyanin emission peak (~ 655 nm).

### Photosystem II activity assay

Oxygen evolution was measured at 20°C in a Clarke-type oxygen electrode (OxyLab2, Hansatech, UK) using cell suspensions at 10 μM Chl, after addition of potassium ferricyanide (Sigma-Aldrich) to 1 mM and the artificial PSII electron acceptor DCBQ (2,6-dichloro-1,4-benzoquinone, Sigma-Aldrich) to 2 mM. Potassium ferricyanide was added from a 200 mM stock solution in water and DCBQ from a 100 mM stock in ethanol. Illumination was with saturating white light (> 20,000 μmol photons m^−2^ s^−1^). Initial rates of O_2_ evolution following the onset of illumination were measured using the OxyTrace+ software from Hansatech. Oxygen evolution in the presence of oxidized DCBQ and saturating light is a direct indicator of the level of PSII reaction centers with functioning water-oxidizing complexes^82^.

## Data availability

The source data underlying Fig. 1b, 1c, 3b-3e, 4, 5b, 5d, 5f, and 5h, in addition to Supplementary Fig. 1b, 1c, 6b, 6c, 9, 10, 11, 12, 13, 14, and 15 are provided as a Source Data file. Microscopy data is available on request. Mass spectrometry data used for Fig. 3 and Supplementary Data 1 were deposited to the ProteomXhange Consortium via PRIDE partner repository with the Project accession PXD019731 (https://www.ebi.ac.uk/pride/archive/projects/PXD019731).

## Acknowledgements

We thank Jennifer Adcott from the Liverpool Centre for Cell Imaging for technical assistance and provision in confocal imaging, and data analysis. We thank Alison Beckett from the Biomedical Electron Microscopy Unit for technical support of electron microscopy imaging and sample preparation. We thank Dr. Hualing Mi for kindly providing the anti-NdhV antibody. This work was supported by the Biotechnology and Biological Sciences Research Council Grants (BB/R003890/1, BB/M024202/1, BB/R01390X/1, L.-N.L.; BB/S003339/1, P.Z; BB/R00370X/1, C.W.M.; BB/R003211/1, P.J.N.), the Royal Society (URF\R\180030, UF120411, RGF\EA\181061, RGF\EA\180233, L.-N.L.), the UK Wellcome Trust Investigator Award (206422/Z/17/Z, P.Z.), the Leverhulme Trust (RPG-2020-054, F.D.C.) and the National Natural Science Foundation of China (32070109, L.-N.L.). We acknowledge Diamond for access and support of the CryoEM facilities at the UK national electron bio-imaging centre (eBIC, proposal NT21004), funded by the Wellcome Trust, MRC and BBSRC.

## Author contributions

T.H. and L.-N.L. conceived the project; T.H., N.T., D.M.S., G.F.D., F.D.C., C.W.M., P.Z., and L.-N.L. performed the research; T.H., N.T., D.M.S., G.F.D., P.B., R.B., P.J.N, C.W.M., P.Z., and L.-N.L. analyzed the data; T.H., N.T., D.M.S., P.Z., and L.-N.L. wrote the manuscript, with the contributions from all the authors. All the authors discussed and commented on the results and the manuscript.

## Supplementary Information

**Supplementary Figure 1.**
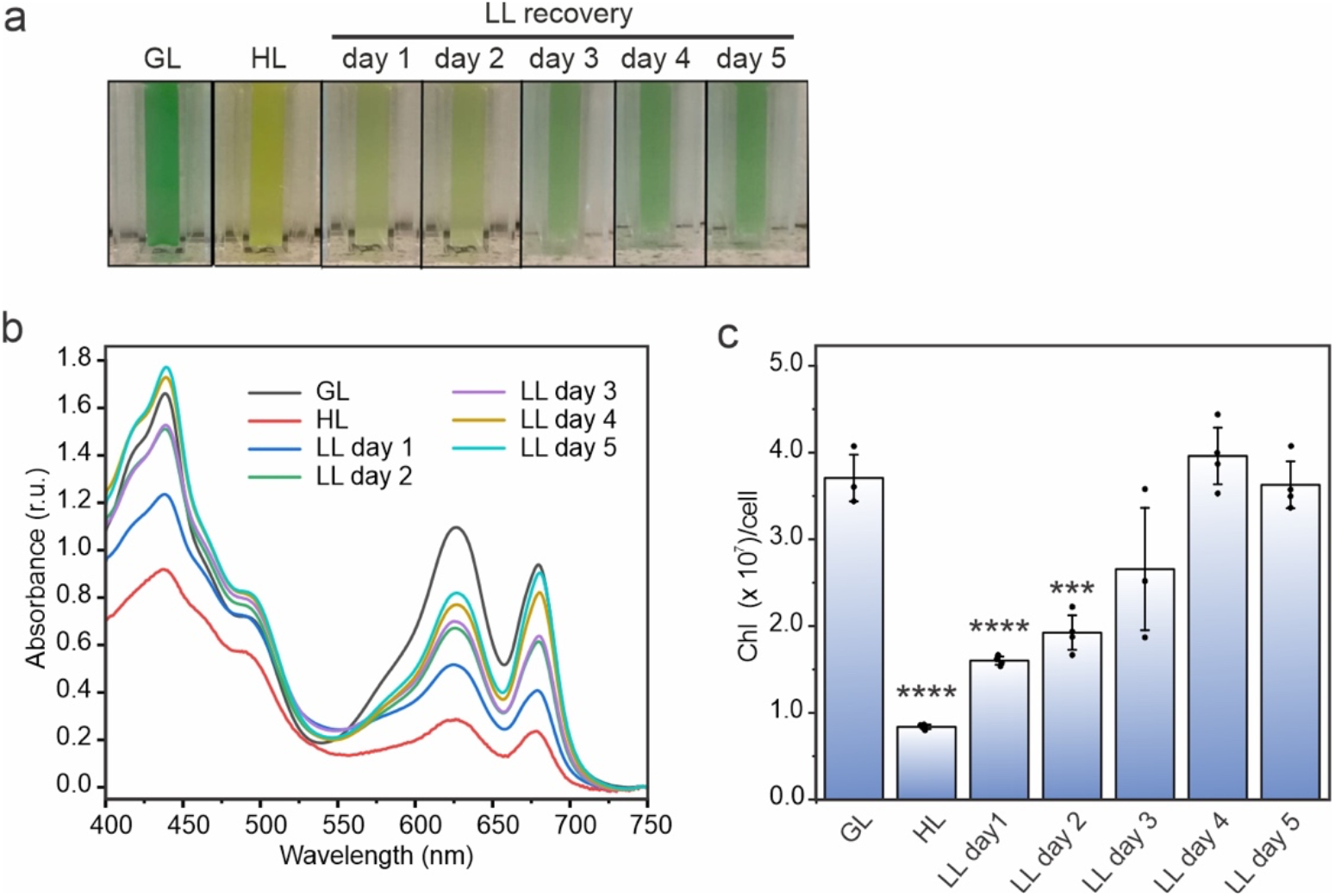
The pigment composition during light-regulated thylakoid membrane biogenesis in *Synechococcus*. Cells were grown under growth light (GL), high light (HL) and HL-grown cells were transferred to low light (LL) conditions for 5 days. **a**, Representative color phenotypes of cells from 3 biologically independent experiments. OD_750_ was adjusted to 0.6 before imaging. **b**, The whole-cell absorption spectra at room temperature. *n* = 3 biologically independent preparations for GL and LL day 3; *n* = 4 biologically independent preparations for HL, LL day 1, LL day 2, LL day 4 and LL day 5. Curves were normalized to 750 nm. **c**, The Chl amount (Chl molecules × 10^7^ per cell). Values are means ± SD; *n* = 3 biologically independent preparations for GL and LL day 3, and *n* = 4 biologically independent preparations for HL, LL day 1, LL day 2, LL day 4 and LL day 5. Asterisks indicate the statistically significant difference compared to GL cells. For HL *p* = 1.0 × 10^−5^, for LL day 1 *p* = 4.89 × 10^−5^ and for LL day 2 *p* = 3.68 × 10^−4^. Statistical analysis was performed using two-sided two-sample t-Test.

**Supplementary Figure 2.**
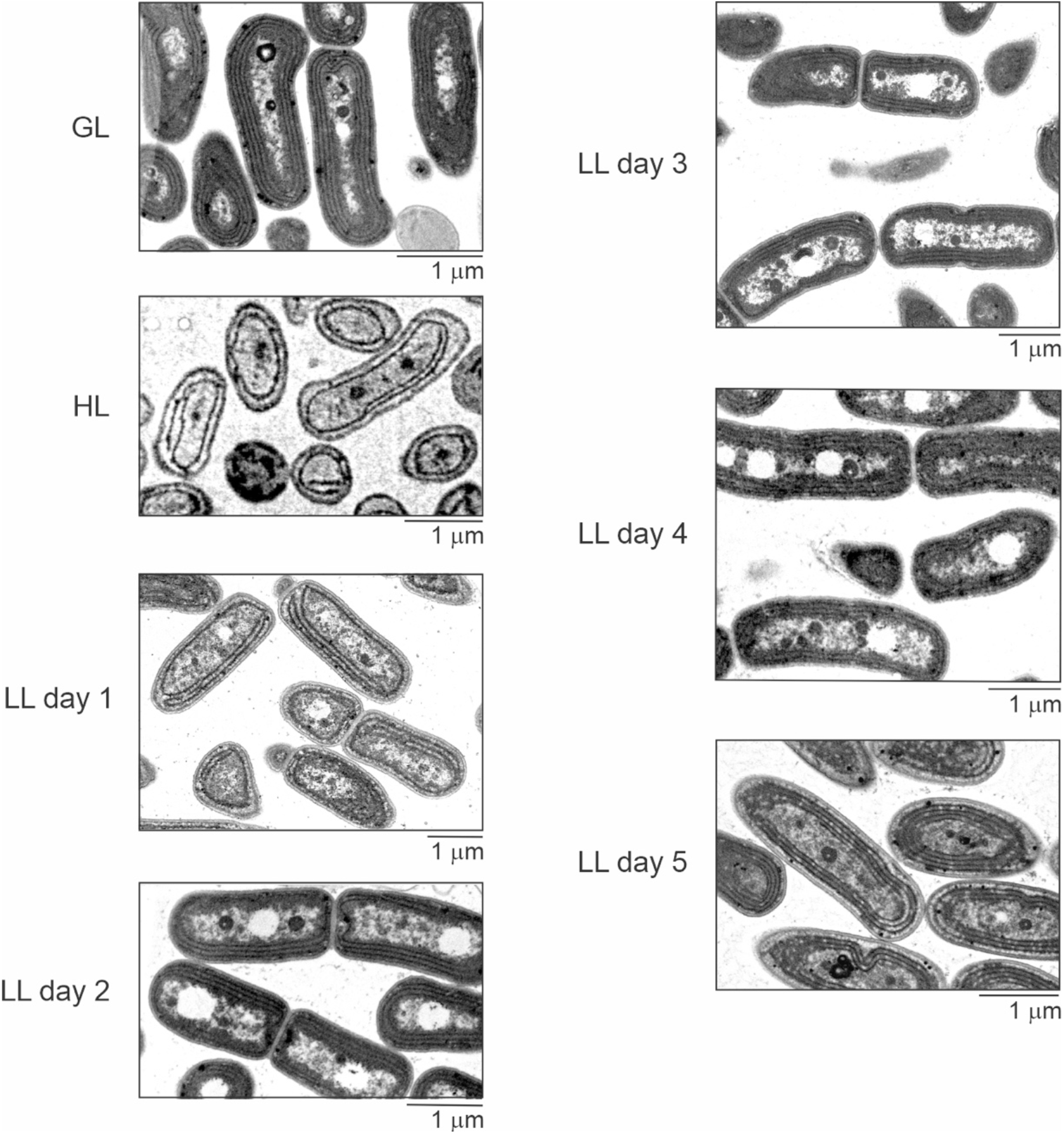
Thin-section transmission electron microscopy during light-regulated thylakoid membrane biogenesis in *Synechococcus*. Cells were grown under growth light (GL), high light (HL) and HL-grown cells were transferred to low light (LL) conditions for 5 days. See also Fig. 1a. Representative TEM images were derived from at least three biologically independent preparations with similar results.

**Supplementary Figure 3.**
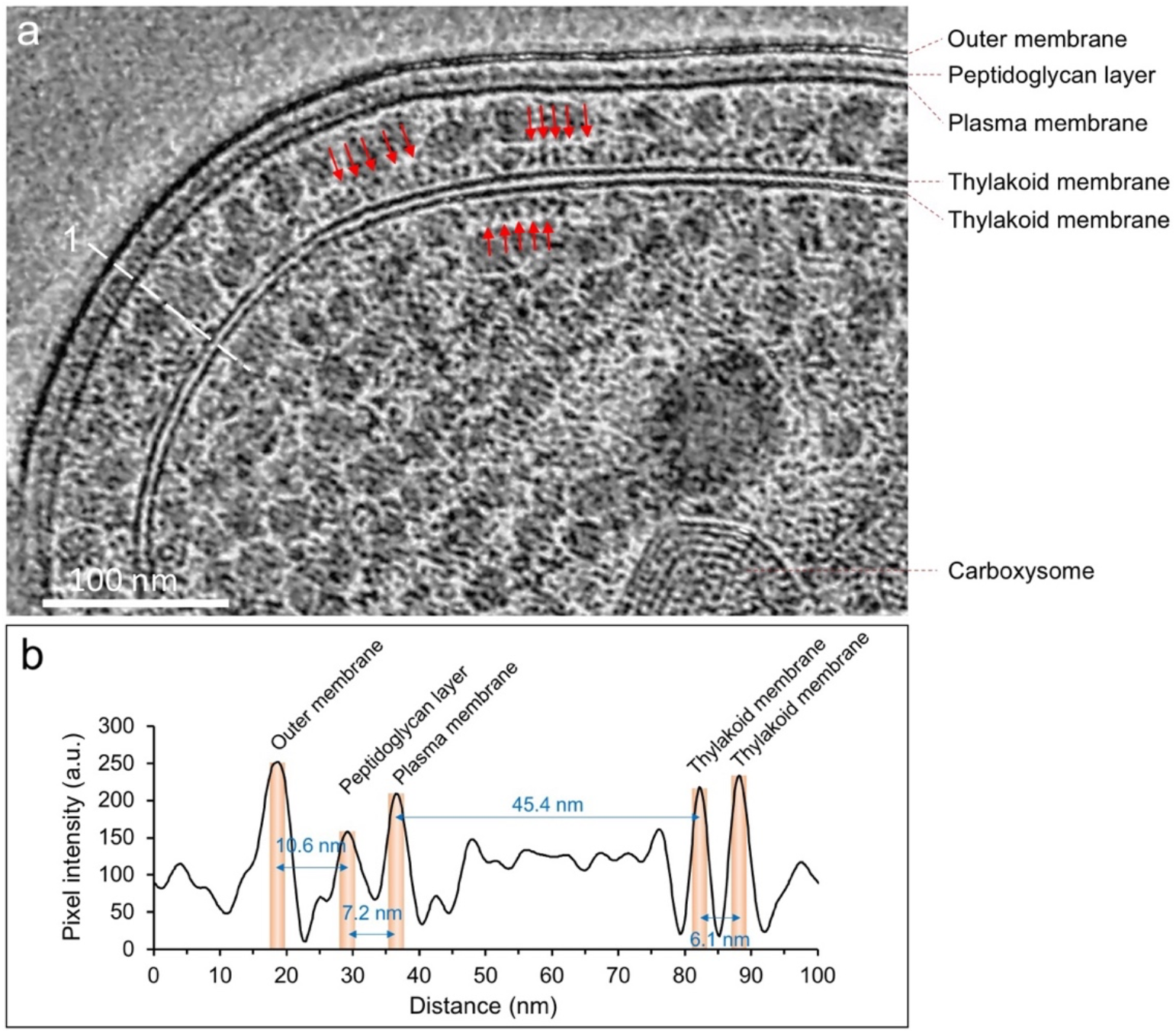
Analysis of the cryo-ET of the *Synechococcus* cell grown under HL. **a**, *in situ* cryo-ET of the *Synechococcus* cell shown in Fig. 2A. Red arrows indicate the phycobilisomes that are densely arranged on the cytoplasmic surface of thylakoid membranes. A line scan (1, line width = 10 pixels) across the outer membrane and thylakoid membranes was used to measure the distances between cellular membranes. **b**, cross-section profile from the line scan indicated in **a**. The outer membrane, peptidoglycan layer, plasma membrane and thylakoid membranes are indicated. a.u., arbitrary units. Profile analysis was performed with Fiji (ImageJ, NIH). The distances between each cellular layer are indicated.

**Supplementary Figure 4.**
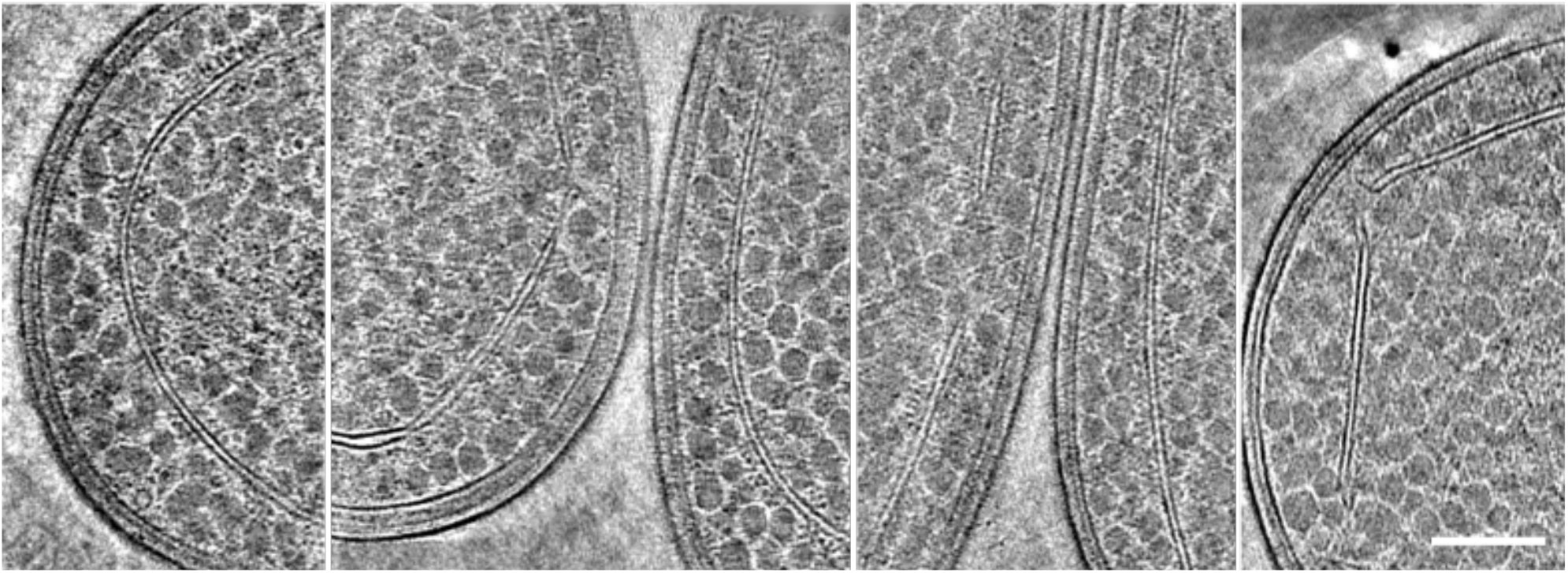
Tomographic slices of representative thylakoid membranes of *Synechococcus*, showing no connections between thylakoid and plasma membranes. Slice thickness, 2.49 nm. Scale bar, 200 nm. Experiment was repeated 3 times with similar results.

**Supplementary Figure 5.**
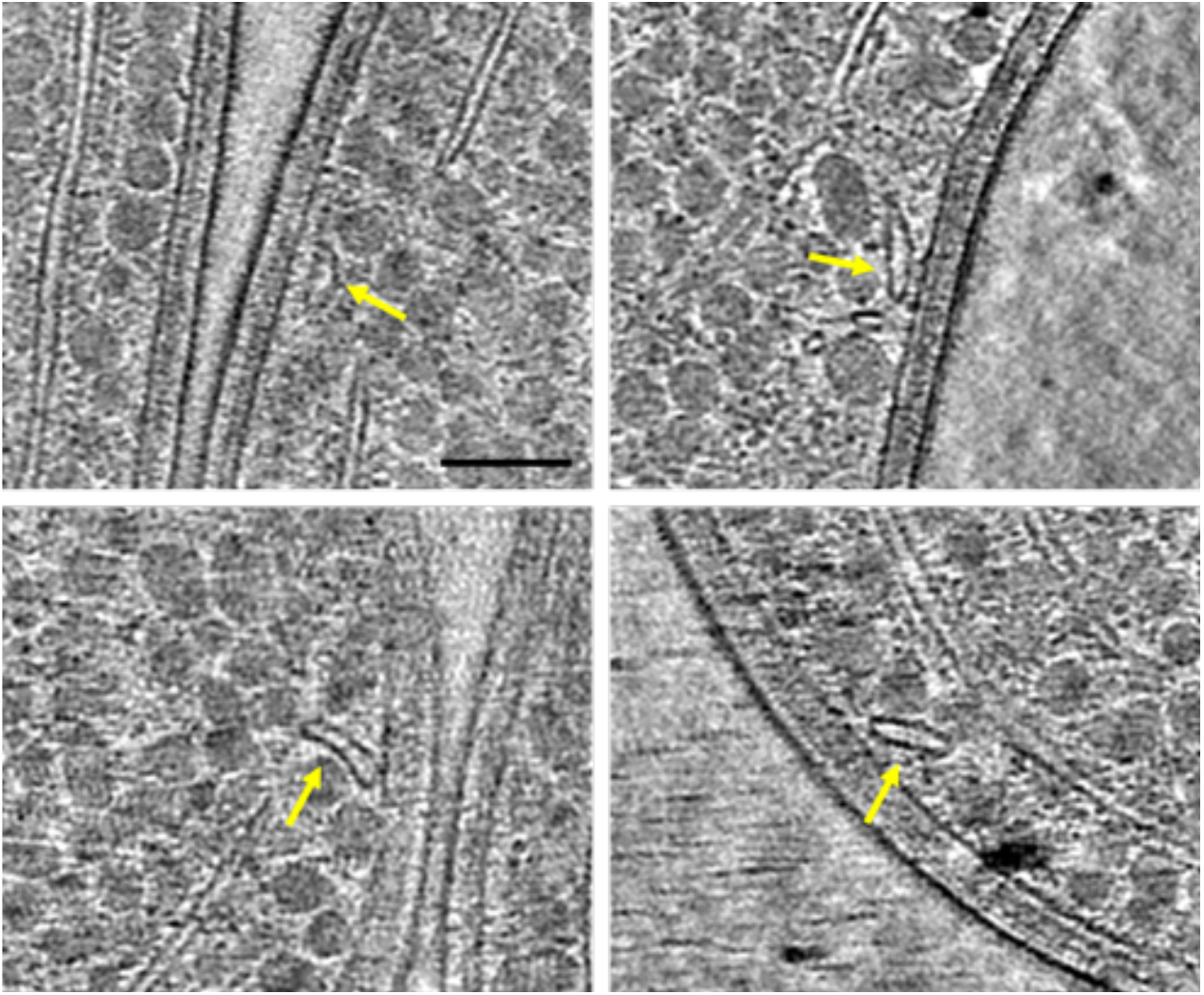
A gallery of small segments of thylakoid membranes in *Synechococcus* near the thylakoid membrane breakages close to the plasma membrane (yellow arrows). Slice thickness, 2.49 nm. Scale bar, 100 nm. See Supplementary Movies 3, 4. Experiment was repeated 3 times with similar results.

**Supplementary Figure 6.**
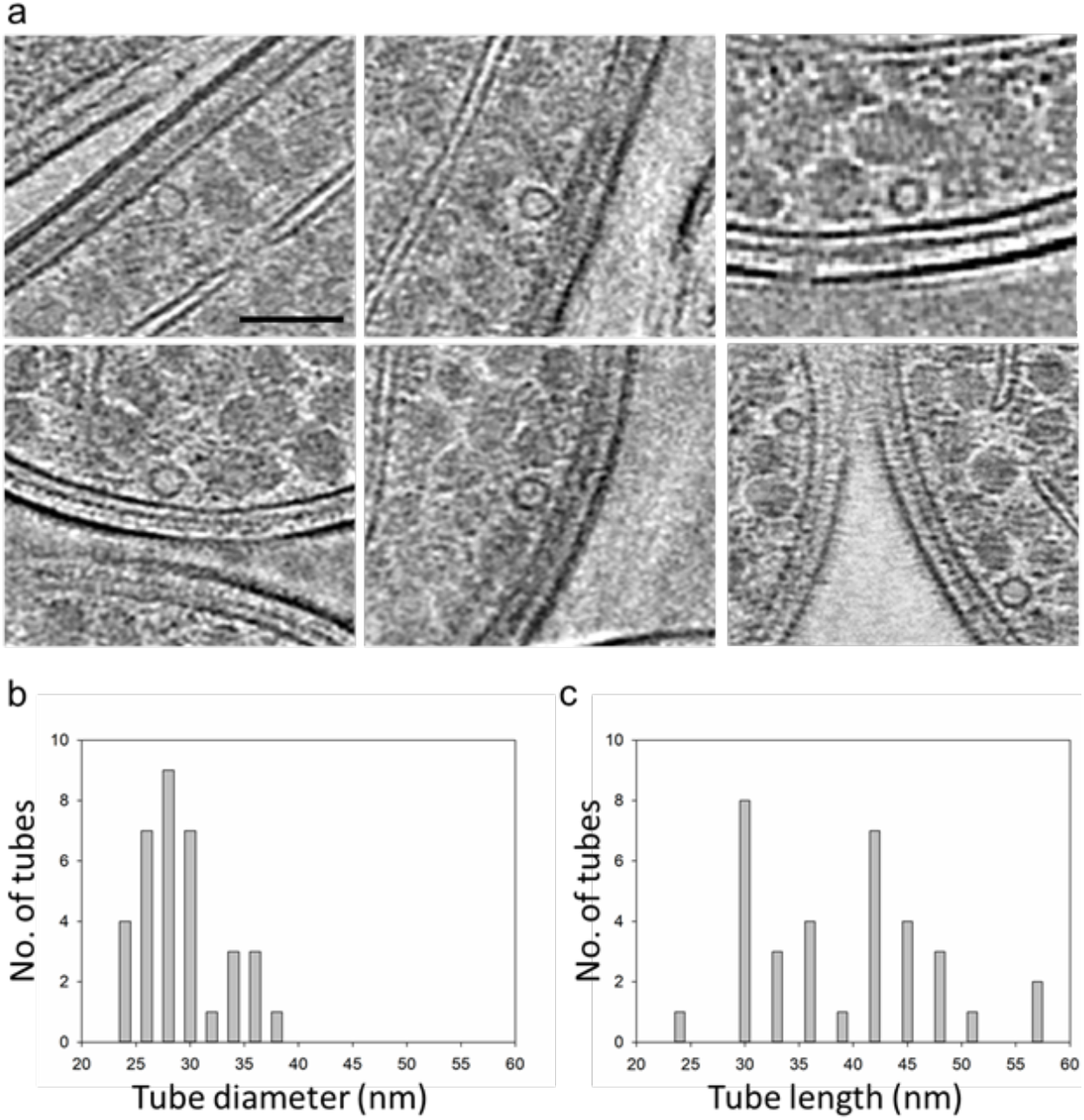
Morphology of the tubular structures *in Synechococcus*. **a**, A gallery of short tubular structures located beneath the cell membrane in tomographic reconstructions. Experiment was repeated 3 times with similar results. **b**, Distribution of the diameter of the tubular structures, *n* = 35 tubular structures. **c**, Distribution of the length of the tubular structures, *n* = 35 tubular structures. Slice thickness, 2.49 nm. Scale bar, 100 nm.

**Supplementary Figure 7.**
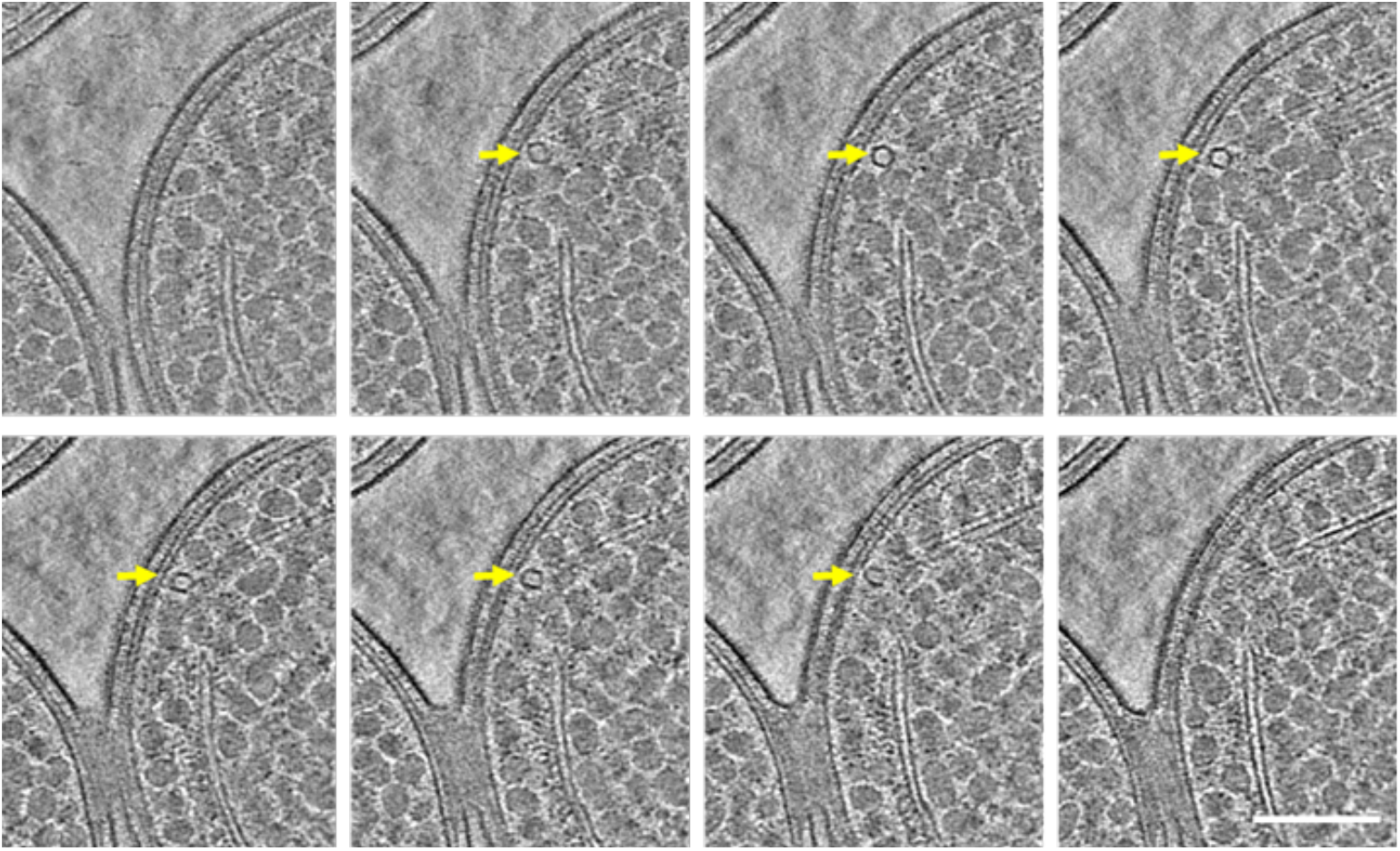
Tomographic slices of a representative small tubular structure close to the thylakoid membrane in *Synechococcus*. Eight evenly spaced single slices (pixel size 2.49 nm), with the distance of 12.5 nm, are shown to enclose a tubular structure. Yellow arrows point to the tubular structure. Scale bar, 200nm. See Supplementary Movie 5. Experiment was repeated 3 times with similar results.

**Supplementary Figure 8.**
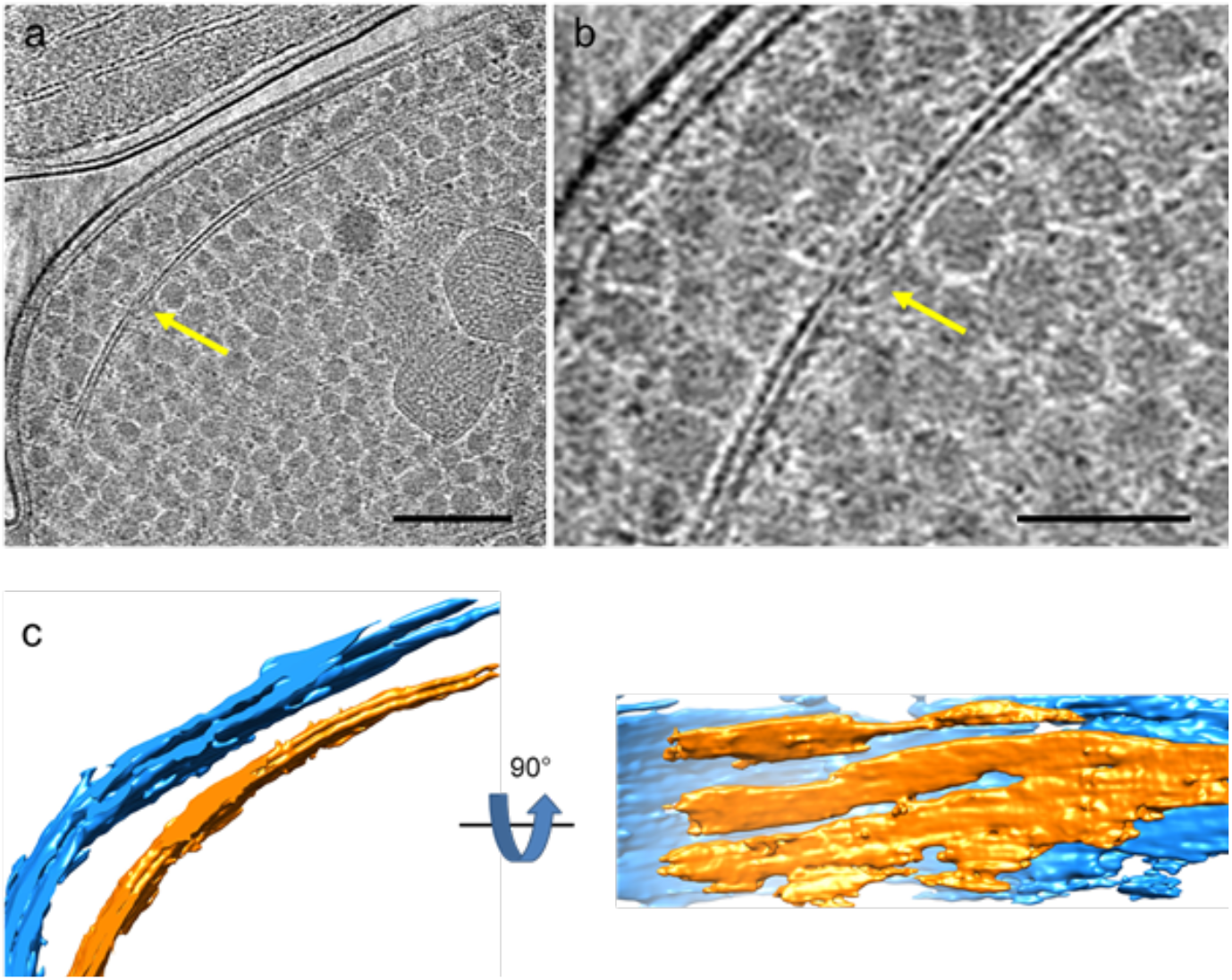
Discontinued thylakoid membranes in *Synechococcus*. **a-b**, Tomographic slice of a *Synechococcus cell* (**a**) and its enlarged view (**b**), displaying perforations of thylakoid membranes (yellow arrows). Experiment was repeated 3 times with similar results. Slice thickness, 2.49 nm. Scale bars, 200 nm in **a** and 100 nm in **b. c**, Segmented volume of the tomogram shown in **a**. Outer and plasma membranes are presented in blue, thylakoid membranes in gold. See Supplementary Movie 6.

**Supplementary Figure 9.**
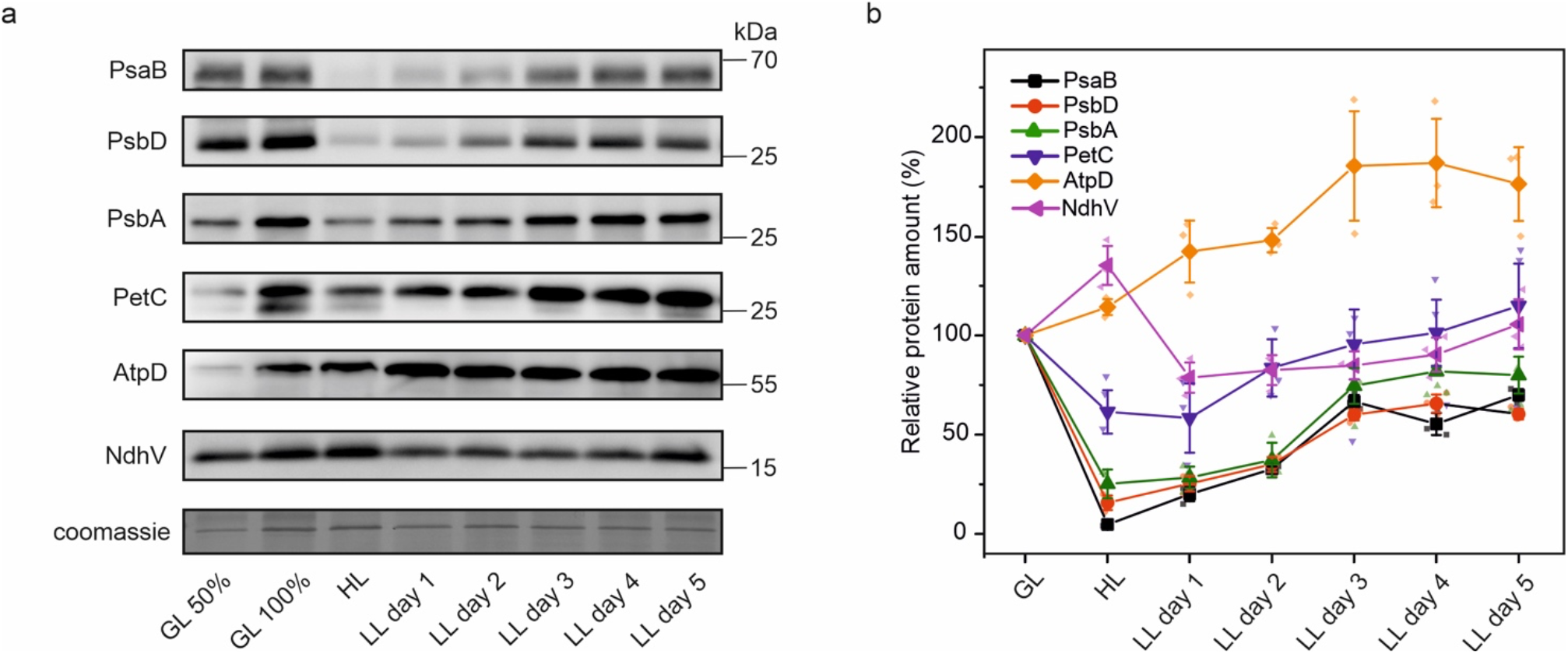
Immunoblot analysis of photosynthetic thylakoid membrane during thylakoid biogenesis. Membrane protein fractions were isolated from *Synechococcus* cells grown under GL, HL, and HL-grown cells transferred to LL. **a**, Immunoblotting with protein-specific antibodies. 10–25 µg (100 %) of isolated membrane proteins were loaded in each lane (15 µg for the immunoblot analysis using α-PsaB, α-PsbD and α-PsbA, 25 µg for α-PetC, α-AtpD and α-NdhV). Coomassie staining is presented as a loading control. **b**, Relative quantification of protein amounts from immunoblots. Values are means ± SD; *n* = 3 biologically independent experiments.

**Supplementary Figure 10.**
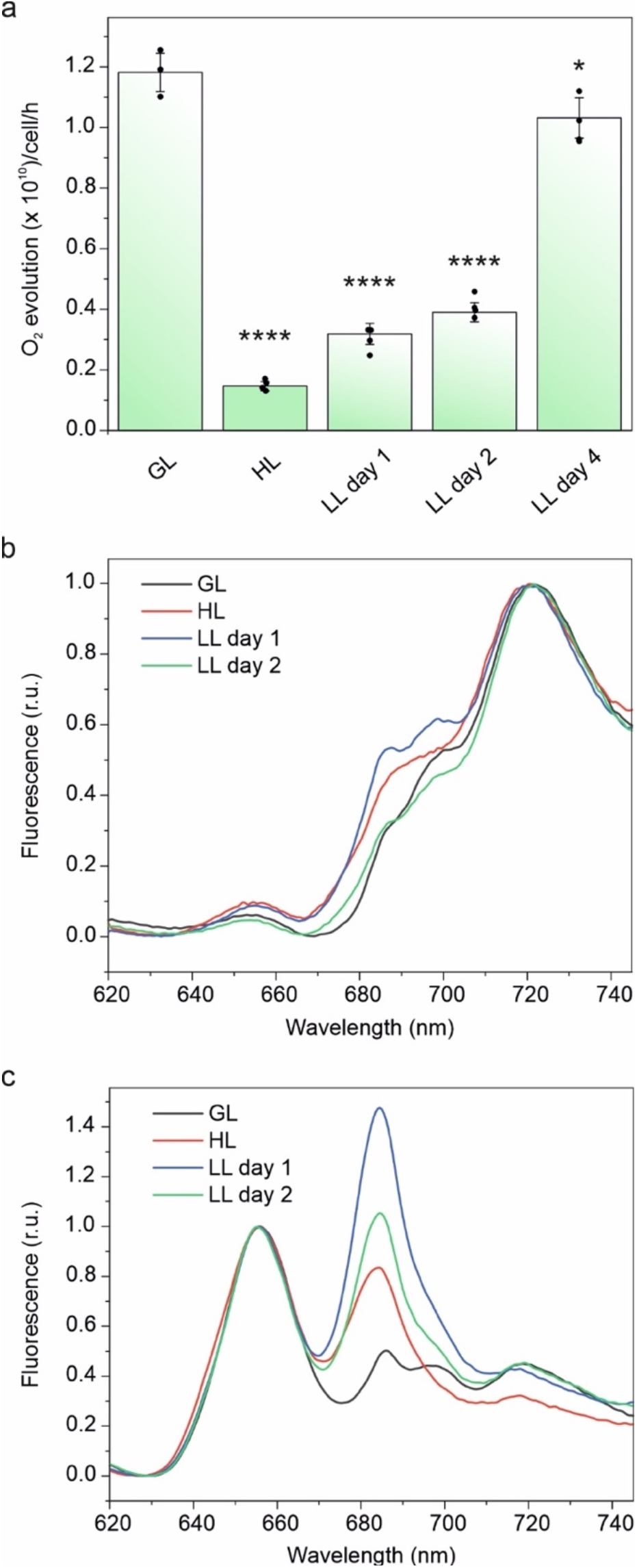
Changes in the functional properties of photosynthetic machinery during light-regulated thylakoid membrane biogenesis in *Synechococcus*. Cells were grown under growth light (GL), high light (HL) and HL-grown cells were transferred to low light (LL) conditions for 2 to 4 days. **a**, The maximum capacity of PSII per cell determined with oxygen electrode in the presence of 2,5-Dichloro-1,4-benzoquinone (DCBQ) and ferricyanide, at saturating light intensity. Values are means ± SD; *n* = 3 biologically independent experiments for GL and *n* = 4 biologically independent experiments for HL, LL day 1, LL day 2 and LL day 4. Asterisks indicate the statistically significant difference compared to GL cells. For HL *p* = 1.41 × 10^−6^, for LL day 1 *p* = 5.88 × 10^−6^, for LL day 2 *p* = 9.88 × 10^−6^ and for LL day 4 *p* = 3.53 × 10^−2^. Statistical analysis was performed using two-sided two-sample t-Test. **b-c**, 77K fluorescence emission spectra when cells were excited with a 435 nm light and were normalized at 720 nm (**b**) or with a 600 nm light and were normalized at 655 nm (**c**). For GL curve is an average of 3biologically independent experiments and for HL, LL day 1 and LL day 2 curves are averages of 4 biologically independent experiments.

**Supplementary Figure 11.**
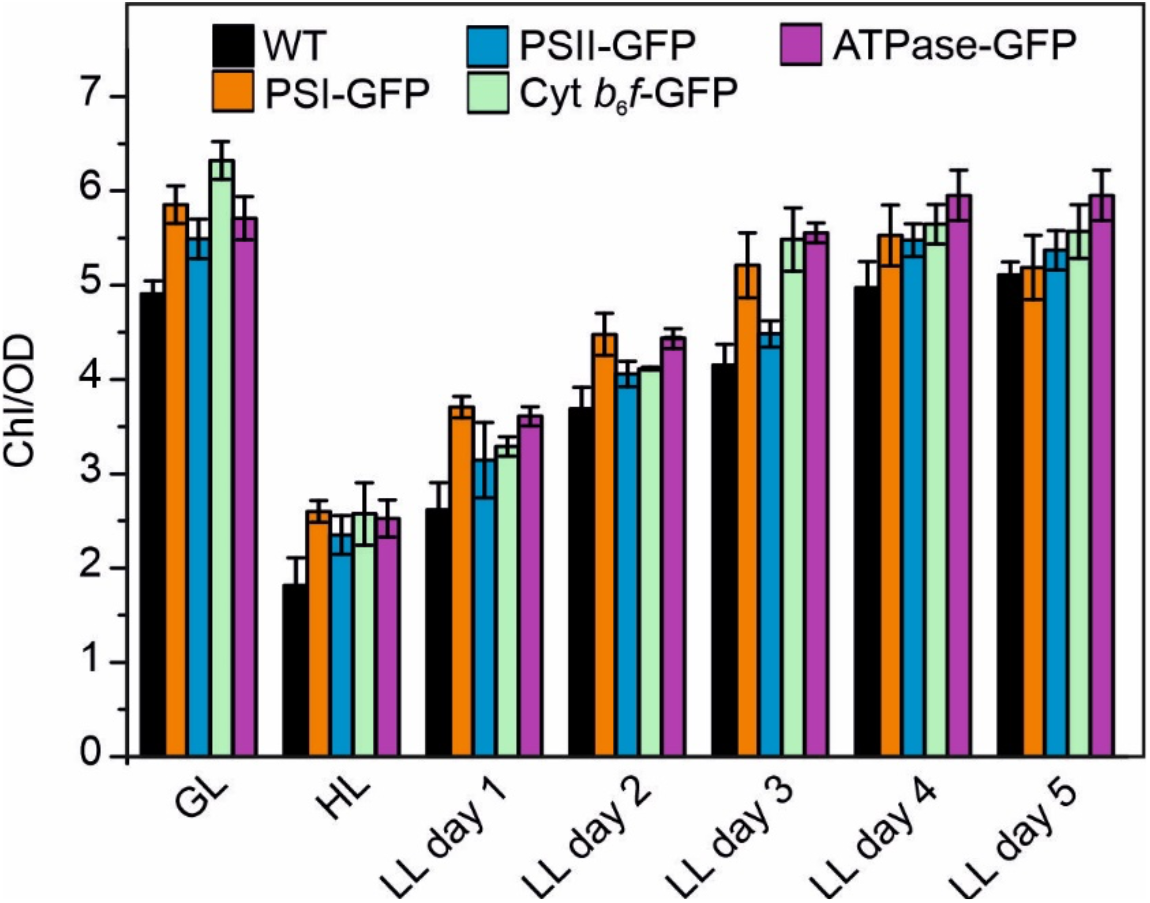
The Chl amount (mg Chl per OD_750_) of WT, PSI-, PSII-, Cyt *b*_6_*f*-, and ATPase-GFP *Synechococcus* strains grown under GL, HL, and HL-grown cells transferred to LL. Values are means ± SD; *n* = 3 biologically independent experiments.

**Supplementary Figure 12.**
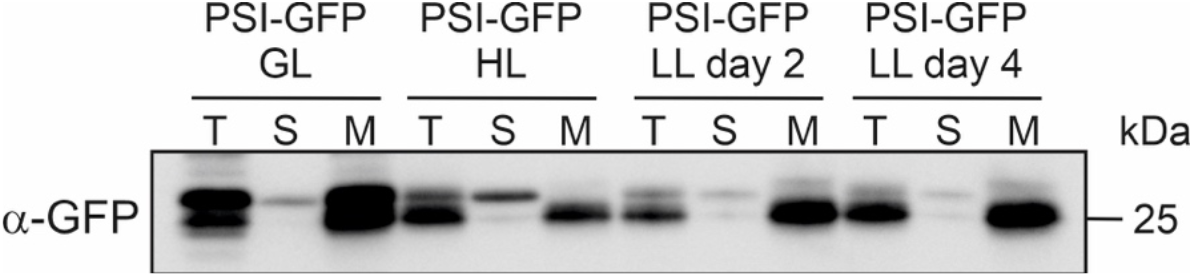
Immunoblotting with GFP-specific antibody from the isolated total protein (T), the soluble (S) the membrane (M) fractions of the *Synechococcus* PSI-eGFP strain grown under GL, HL, and HL-grown cells transferred to LL for 2 and 4 days. 30 µg of proteins from each isolated fraction were loaded in each lane. Experiment was repeated 3 times with similar results.

**Supplemental Figure 13.**
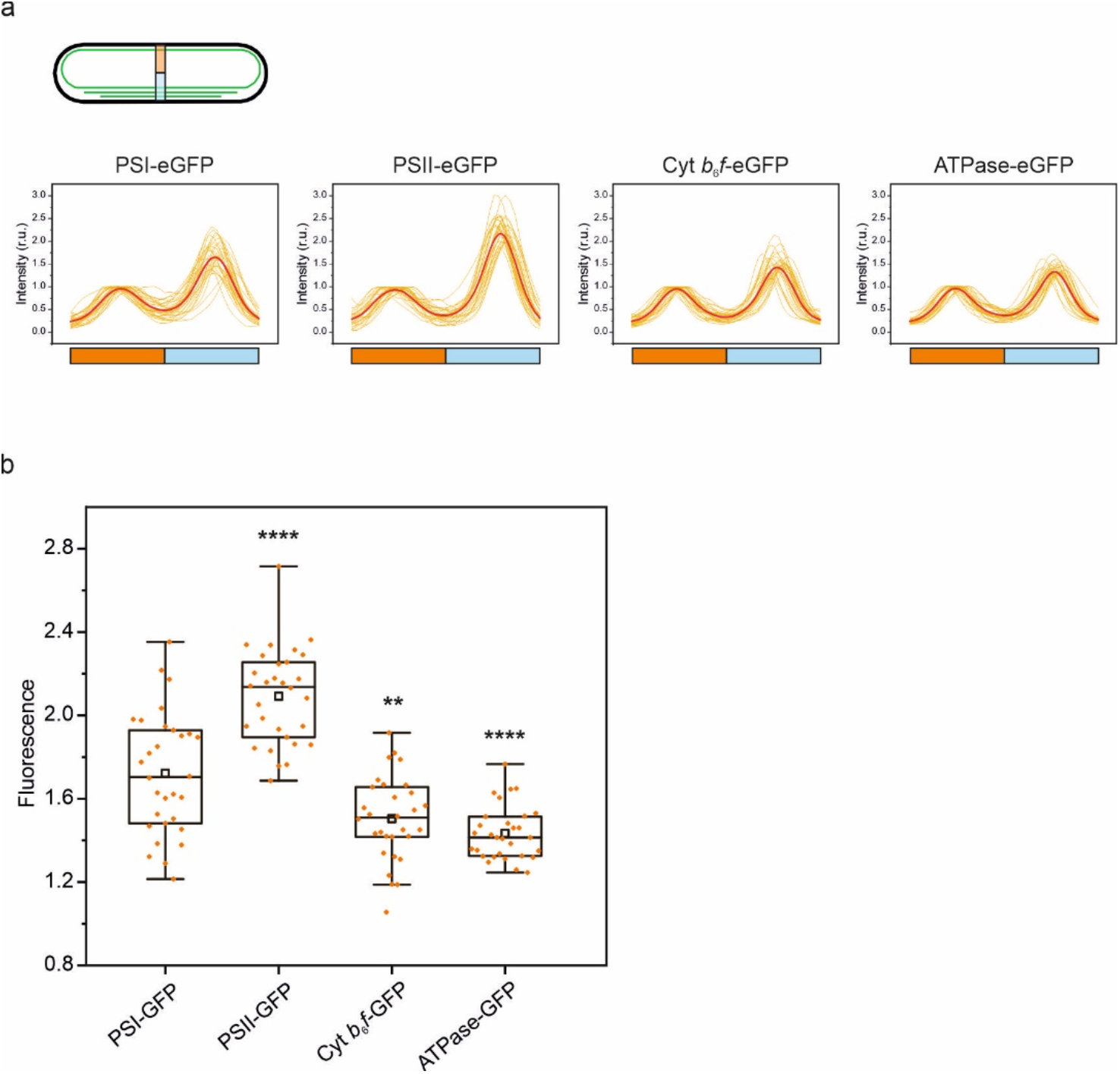
Distribution of photosynthetic protein complexes between longitudinal cell sides at the beginning of thylakoid membrane biogenesis. **a**, Intensity of PSI-, PSII-, Cyt *b*_6_*f*- and ATPase-eGFP signal across the cell in *Synechococcus* strains transferred from HL to LL conditions for one day. Profiles in red are averages from 30 individual cells (in yellow on the background) from 3 biologically independent experiments. The PSI-GFP, PSII-GFP, Cyt *b*_6_*f*-GFP, and ATPase-GFP signals displayed a similar asymmetrical distribution as Chl autofluorescence signals. Orange bar in X-axis represents the longitudinal cell side with weaker Chl fluorescence representing fewer thylakoid layers (GFP fluorescence was normalized to 1), light blue bar with stronger Chl fluorescence representing more thylakoid layers. **b**, The unnormalized signal ratios of PSI-, PSII-, and Cyt *b*_6_*f*-eGFP between longitudinal cell sides. *n* = 30 cells from three biologically independent experiments. The higher value corresponds to more uneven distribution between longitudinal cell sides. Box plots display the median (line), the average (open square), the interquartile range (box) and the whiskers (extending 1.5 times the interquartile range). Asterisks indicate the statistically significant differences compared to PSI-GFP signal. For PSII-GFP *p* = 1.08 × 10^−6^, for Cyt *b*_6_*f*-eGFP *p* = 1.37 × 10^−3^, and for ATPase-GFP *p* = 6.56 × 10^−6^. Statistical analysis was performed using two-sided two-sample t-Test.

**Supplementary Figure 14.**
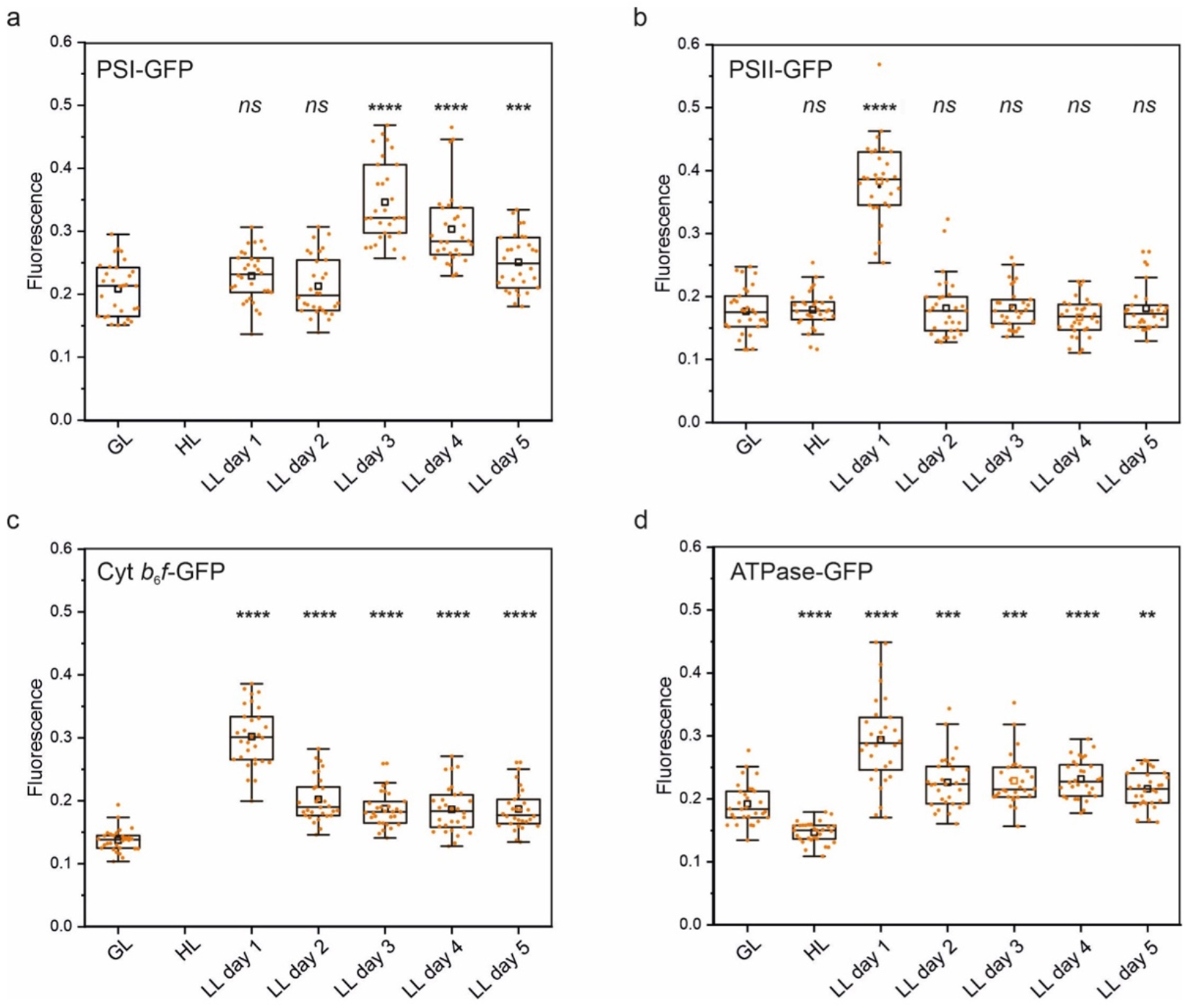
Normalized deviation of eGFP-signal distribution along the thylakoid membranes in the *Synechococcus* strains grown under GL, HL, and HL-grown cells transferred to LL for 5 days. **a**, PSI-GFP, **b**, PSII-GFP, **c**, Cyt *b*_6_*f*-GFP, and **d**, ATPase-GFP. *n* = 30 cells from 3 biologically independent experiments. The standard deviation of the signal normalized to the total fluorescence provides a quantitative measure of the patchiness of the signal. Evenly distributed fluorescence fluctuates little along the line profile and therefore has a low standard deviation, whereas the patchy distribution has a high standard deviation. Box plots display the median (line), the average (open square), the interquartile range (box) and the whiskers (extending 1.5 times the interquartile range). Asterisks indicate statistically significant difference to GL cells. For PSI-GFP LL3 *p* = 3.37 × 10^−14^, for PSI-GFP LL3 *p* = 2.32 × 10^−9^, for PSII-GFP LL 1 *p* = 2.65 × 10^−22^, for Cyt *b*_6_*f*-GFP LL1 *p* = 4.17 × 10^−25^, for Cyt *b*_6_*f*-GFP LL2 *p* = 1.52 × 10^−12^, for Cyt *b*_6_*f*-GFP LL3 *p* = 6.11 × 10^−11^, for Cyt *b*_6_*f*-GFP LL4 *p* = 2.42 × 10^−8^, for Cyt *b*_6_*f*-GFP LL5 *p* = 1.94 × 10^−9^, for ATPase GFP HL *p* = 5.97 × 10^−9^, for ATPase GFP LL1 *p* = 1.88 × 10^−9^, for ATPase GFP LL2 *p* = 7.22 × 10^−4^, for ATPase GFP LL3 *p* = 2.45 × 10^−4^, for ATPase GFP LL4 *p* = 8.97 × 10^−6^ and for ATPase GFP LL5 *p* = 3.35 × 10^−4^. *ns*, not significant. Statistical analysis was performed using two-sided two-sample t-Test. Under HL PSI- and Cyt *b*_6_*f*-GFP signals were not quantified due to unrestricted and weak signals, respectively.

**Supplementary Figure 15.**
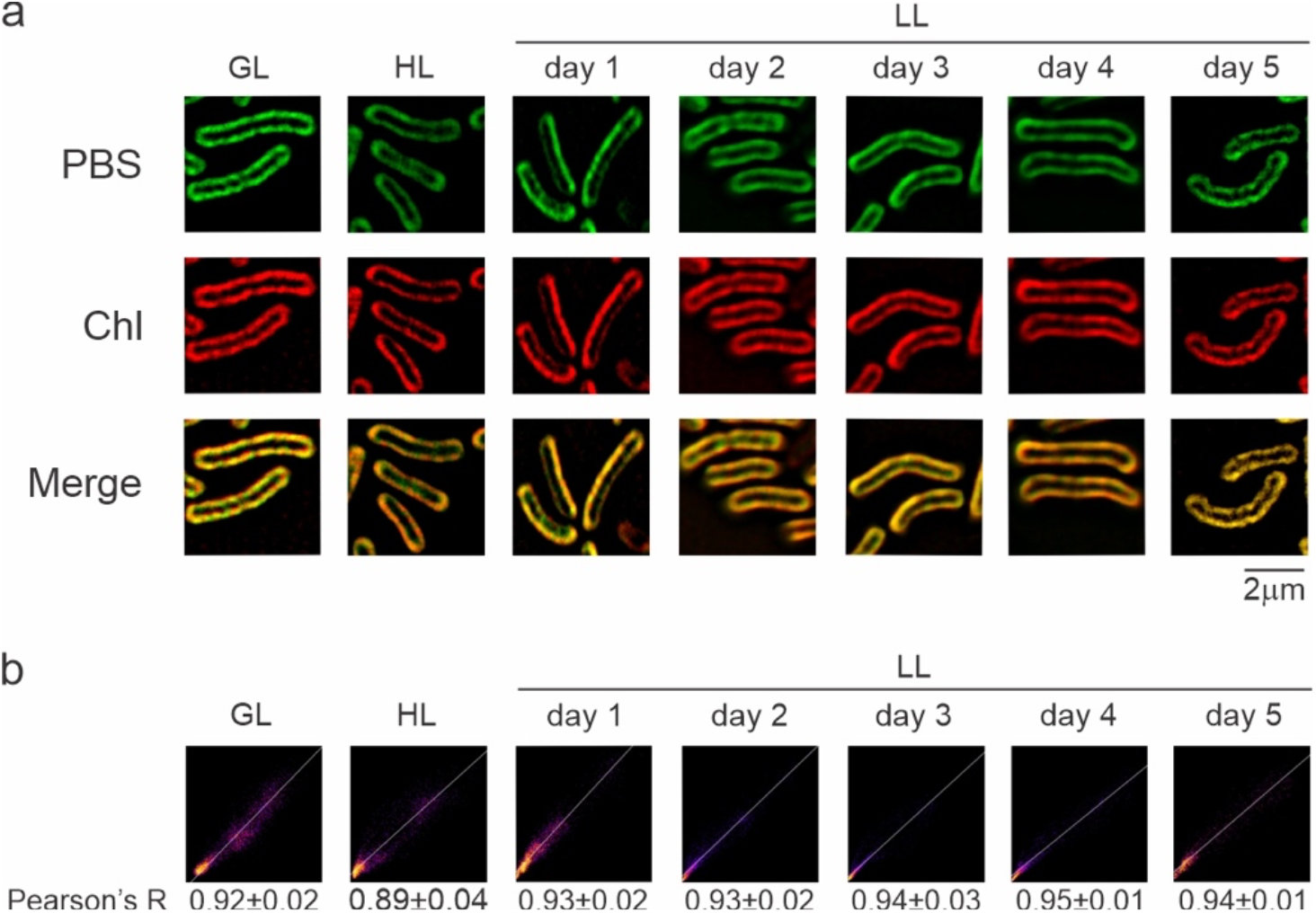
Expression and intracellular localization of phycobilisomes (PBS) *in vivo* in *Synechococcus* during thylakoid membrane biogenesis. **a**, *Synechococcus* wild type (WT) cells imaged by Dragonfly spinning disk confocal microscope with the super-resolution radial fluctuations (SRRF)-stream technology. Figures are representative of 3 biologically independent experiments. First row: phycobilisome (PBS) fluorescence; second row: Chl autofluorescence; third row: merged channels. **b**, Scatter plots and Pearson’s correlation values (± SD) for the colocalization of PBS and Chl for each studied condition. PBS and Chl fluorescence were analyzed from 30 individual cells in total from 3 biologically independent experiments.

**Supplementary Table 1.**
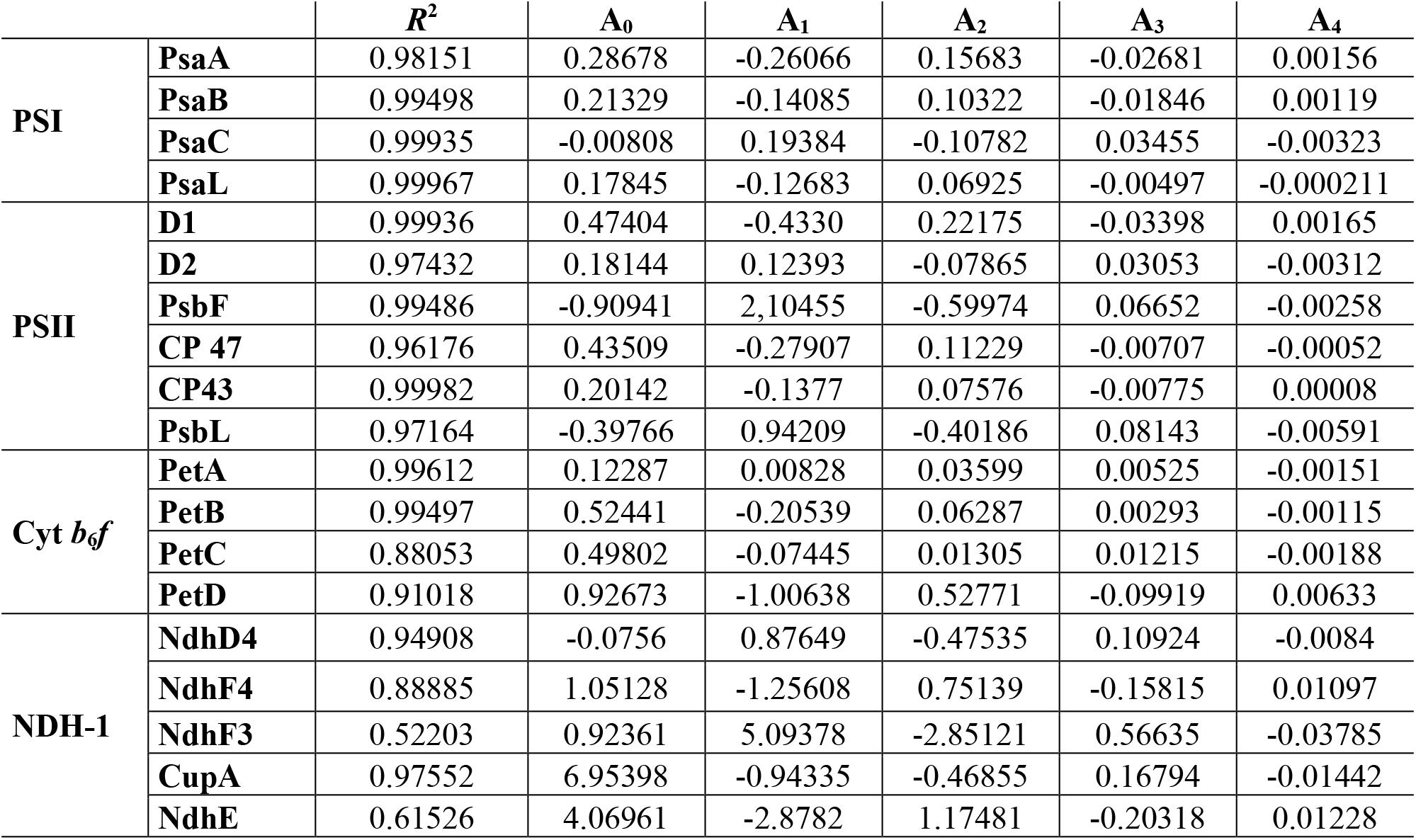
Adjusted *R*^2^ values and coefficient factors of the fourth-degree polynomial functions (y = A_0_ + A_1_×x + A_2_×x^2^ + A_3_×x^3^ + A_4_×x^4^) fitted for representative photosynthetic proteins from global quantification as a function of time.

